# Universal Baseline for *in vitro* Selection of Genetically Encoded Libraries

**DOI:** 10.64898/2026.02.14.705946

**Authors:** Kejia Yan, Guilherme M. Lima, Tara Bahadur, Vincent Albert, Zoe O’Gara, Gary Bao, Christin Kossmann, William Kirby, Fernando B. Mejia, Matthew L. Michnik, Kristen Maiorana, Ratmir Derda

## Abstract

Genetically encoded (GE) libraries enable identification of high-affinity ligands for diverse molecular targets through iterative *in vitro* selection and DNA sequencing or next-generation sequencing (NGS). Despite their impact in therapeutic development, a systematic framework for evaluating reproducibility in GE-molecular discoveries remains limited. To aid such analysis, we introduce the concept of baseline response, which reproducibly partitions active and inactive members of *in vitro* selection. The baseline response is provided by spiking a random DNA-barcoded population. We calibrated the baseline concept using Bioconductor EdgeR differential enrichment (DE) analysis of NGS of phage-displayed selection on oligosaccharide chitin and hepatitis virus NS3a* protease as model targets. We further show that mixing discovery campaigns also offers an effective baseline: chitin-enriched peptides serve as a baseline for DE-analysis of NS3a* selection and NS3a*-enriched peptides serve as a baseline for chitin binders. We applied baseline-stratified DE-analysis to 66 parallel selections performed in 3–5 replicates across 22 extracellular targets, including HER1-3, EpCAM, CAIX, PD-L1, and eight integrin receptors. Automated DE-analysis across hundreds of NGS files produced hits validated in a secondary screen and yielded synthetic macrocyclic ligands with mid-nanomolar affinity confirmed in 2–3 biophysical assays. For PD-L1, we further demonstrated how baseline-calibrated NGS data provide decision-enabling information for optimization of peptide macrocycles to yield potent single-digit nanomolar ligands for the cell-surface receptor. We anticipate that baseline-based analyses of NGS data from *in vitro* selection procedures will offer a scalable framework for reproducible hit discovery and standardized analysis across diverse *in vitro* selection campaigns.

**Significance Statement:** Genetically encoded selection technologies such as phage, mRNA and ribosome display, have produced FDA-approved therapeutics and numerous clinical candidates. Yet reproducibility in such *in vitro* discovery systems is rarely evaluated against a defined experimental baseline. Here, we establish a universal baseline by spiking unrelated, DNA-barcoded peptide sequences into selection libraries and quantifying their binding alongside target-enriched populations. This composition-agnostic strategy enables rigorous normalization, confidence assessment, and cross-target comparison of molecular discovery outcomes. Our framework introduces practical standards for reproducibility and statistical benchmarking across genetically encoded display platforms.

## Introduction

In a global therapeutic market, a substantial fraction of approved therapeutics and clinical candidates undergoing clinical trials originate from molecules discovered in genetically-encoded selection processes (1, 2). Phage display (3) and related display technologies (ribosome, mRNA-, and cell-display) (4–7), aptamer selection (a.k.a. SELEX) (8), and selection from DNA-encoded libraries (DEL) (2, 9) all represent the same process of DNA-encoded molecular discovery performed by iterative *in vitro* selection. This process usually starts from million-to-billion scale diverse libraries of DNA-encoded molecules and aims to enrich a subset of these molecules that bind to a target of interest. Most approaches, with exception of traditional DEL, employ multiple rounds of *in vitro* selection and amplification cycles. At each selection cycle, a collection of encoded molecules is incubated with a target displayed on the surface of the microtiter well plate, bead, or cell surface. This selection is conceptually identical to a traditional binding assay or high-throughput screening (HTS) (10) but fundamental differences exist in biophysical analysis of these processes. Reproducibility and baseline response are fundamental in the analysis of binding assays, the same analysis of reproducibility over a baseline population gives rise to robust statistical analyses of HTS based on binding assays; examples are Z’ score analysis of HTS assays. To allow equivalent assessment of the reproducibility in molecular discovery emanating from *in vitro* selection of DNA-encoded libraries, the field critically lacks a unified concept of discovery baseline which can be universally employed in any molecular discovery powered by *in vitro* selection.

To our knowledge, discussion of reproducibility, baseline and robust statistics derived from thereof are scarce in publications that employ *in vitro* selections: Krusemark and co-workers employed Z’-type analysis to analyze DEL selection (11) and our lab repurposed Z’ analysis to test robustness of selection of phage-displayed glycans (12). Phage display literature evaluate convergent selection to the same target-unrelated peptides (13), or the reproducible discovery of parasitic sequences (14). Common approach to assess robustness of selections in peptide-display and antibody-display literature involves monitoring enrichment of pre-defined binding clones spiked into a naïve library (3), or reproducible emergence of specific motifs, such as streptavidin-binding peptides with HPQ motif (15), integrin-binding peptides with RGD-motif (16–18) or known antigen sequences (19). Similarly, DEL and SELEX publications often benchmark selection procedures by analyzing emergence of hits for clinically-relevant targets (undruggable transcription factors (20), prostate-specific antigens (21), bromodomain (22, 23), carbonic anhydrase (24), thrombin (25), etc.) Several publications explored reproducibility of discoveries as a function of display formats (26) or as a function of the size of mRNA-displayed libraries (27) or phage-displayed libraries (28), or SELEX libraries (29). In the latter example, Szostak and co-workers found relationships between the complexity of the library, robustness of the selection and the information content of the evolved aptamers (30). Convergence of parallel evolution of proteins (31), ribozymes (32), and phenotypic outcomes in in different species (33) have been reported. Notable example from Liu and co-workers quantified reproducible convergence of selection trajectories in continuous phage assisted selections of T7 RNA polymerase (34). These and other analyses rely on observation of specific molecular outcomes, such as reproducible enrichment of defined sequences, molecular structures, or motifs. Recent translation to automation of large NGS datasets emanating from selection procedures and use of such datasets for machine learning (ML) requires robust stratification of active and inactive populations in such datasets while limiting the number of false-positive and false-negative hits (35, 36). An anticipated sequence-specific-outcomes or predefined targets-specific decisions can improve analyses of the selection procedures, but an unbiased analysis of *in vitro* selection can be improved with a universal, composition-agnostic analysis methods. We strived to devise the analysis of the reproducibility of molecular discoveries that does not rely on any analysis of molecular composition or pre-defined knowledge of molecular structure. For example, the analysis of reproducibility in binding assays is independent of the molecular identity of binding partners. Instead, binding assays employ the “baseline response” as a foundation for statistical analysis (37, 38). Agglomeration of binding data stratified by baseline is the foundation of all large-scale analysis of binding data (e.g., HTS), structure activity relationship (S.A.R.) derived from such data and Machine Learning approaches that learn from data (39, 40). In this manuscript, we implemented a universal experimental baseline to quantify and normalize molecular discovery outcomes across multiple targets and multiple *in vitro* selection procedures.

Immunization of animals is an example of a molecular discovery process that employs a universal baseline established nearly half-a-century ago. In discovery of antibody A to an antigen X, baseline response is produced by sampling antibodies from the serum of non-immunized animal and measuring their binding to antigen X (41). From here one can extrapolate that a baseline response **B** for any molecular discovery that employs a mixed molecular library [L] and *in vitro* selection towards target T. Specifically, **B** is binding of collection of molecules randomly sampled from [L] towards target T. By analogy, we demonstrate how to implement such baselines in discoveries that employ randomized library of peptides [P] displayed on phage. While the rest of the manuscript focuses on phage display and peptide libraries, we note that identical principle can be used to define a baseline for any in vitro selection procedure that employs large mixed populations of encoded molecules and iterative multi-round selection procedures.

A *bona fide* random baseline sampled from a native molecular library [L] contains both active and inactive components. A distribution of the affinities in such random naïve samples have been estimated in a hallmark publication by Lancet *et al*. (42) who measured the shape of the distribution in the naïve and selected populations of antibodies by equilibrium dialysis. By definition, *in vitro* selection eliminates the non-fit members of the library. Hence, after several rounds of *in vitro* selections, the baseline responses no longer exist in such selected populations. In this closed population it is often impossible to establish a robust baseline response because all members of this population exhibit a various degree of binding to the desired target. To restore this lost baseline, we spiked a universal DNA-barcoded subset of random peptide sequences into every step of the molecular discovery pipeline. This random baseline population, distinguishable from the remainder of the population through silent codon barcoding (43, 44), delineates a true baseline response from target-specific selections. With this approach, we performed selections across 24 protein targets and observed that the presence of a universal *in situ* baseline enables quantitative assessment of enrichment fidelity, reproducibility and cross-sectional analysis of discoveries performed for different targets and different library compositions. By decoupling selection-specific effects from background noise, our baseline strategy establishes a statistical foundation for benchmarking *in vitro* selections and provides a missing component for reproducibility and data normalization in genetically encoded molecular discovery pipelines.

## Results

### Spiked non-binding baseline into pre-enriched library populations

We tested whether a spiked non-targeted (baseline) library provides any additional information in a traditional selection of phage displayed library. As a model, we utilized a previously characterized selection (45) against hepatitis C virus NS3a variant (NS3a*) immobilized on streptavidin magnetic beads (SMB). NS3a*-bound peptides were eluted using grazoprevir (**Fig. 1A**). In a standard procedure without baseline (**Fig. 1A-B**), hits were identified as peptides significantly enriched in differential enrichment (DE) analysis of NGS data from the third round of selection (R3). The scatter plot of input vs. output NGS reads (**Fig. 1C**) had a characteristic bimodal distribution: the peptides that pass DE analysis reside in the area clearly separated from the peptides that reside on the diagonal. The latter “on-diagonal” population contains peptides that have similar input and output NGS reads. In addition to selection on NS3a* target, we employed “control” selection procedures, here SMB without NS3a*, to delineate the peptides enriched due to association with NS3a* form those that might be binding to SMB. The scatter plot for this control selection (**Fig. 1D**) has the same bimodal shape with clearly separated SMB-enriched “off diagonal” and non-enriched “on diagonal” population. Analysis of Figure 1c-d concludes that populations in “target” and “control” selections appear to be similar with one exception: more peptides reside in the off-diagonal (enriched) population in target selection. Peptides enriched on NS3a* but not SMB contained a recurrent DMT motif, which was previously confirmed to be a hallmark of NS3a*-binding macrocycles.

**Figure 1.**
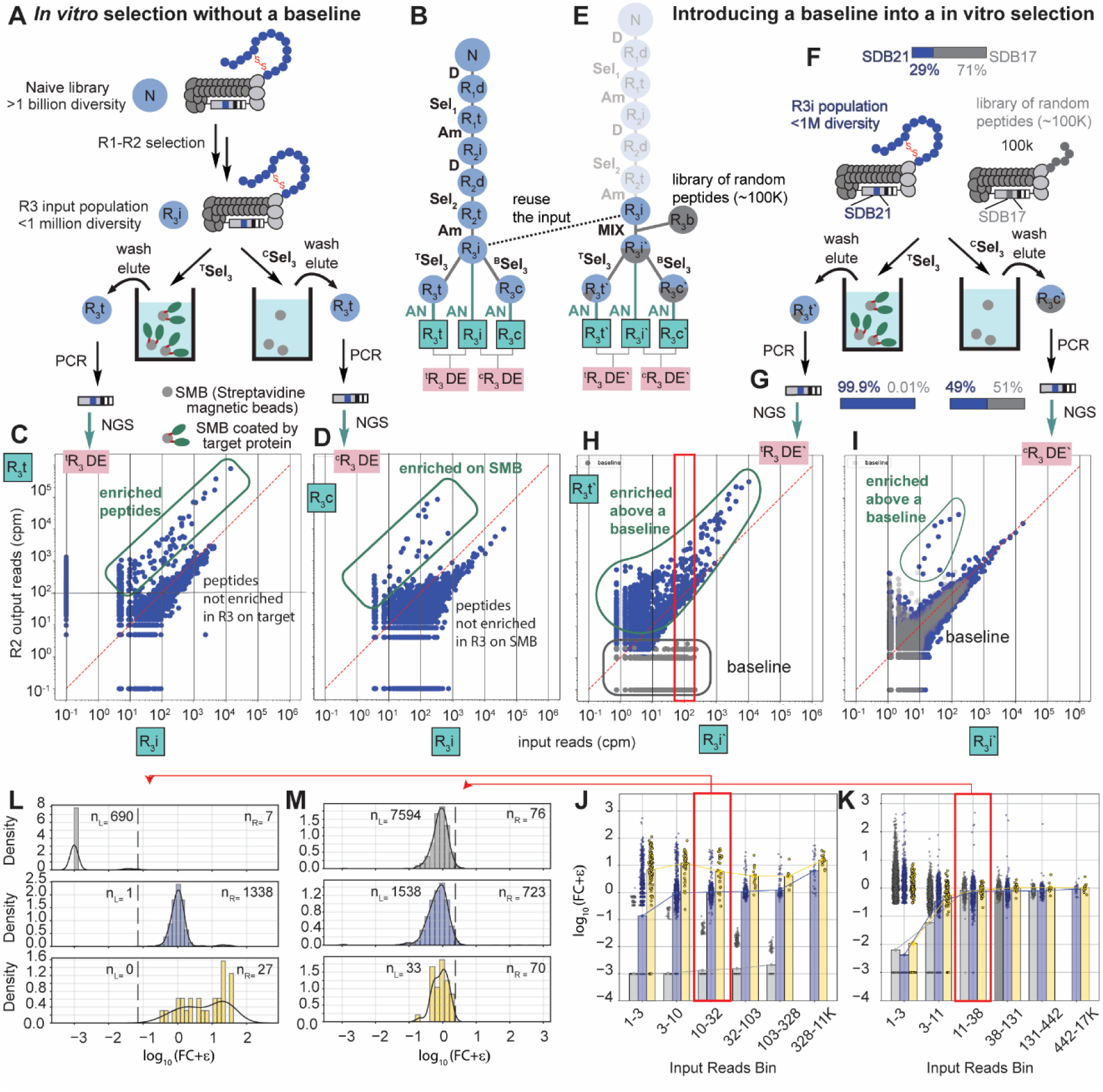
Establishing a baseline for evaluating selection quality using a mixed non-binding control library in the final round of selection. A) Procedure of final round selection without baseline. B) Flow chart of panning campaign against NS3a* without baseline. C) Scatter plot of normalized output versus input read counts against target. Peptides enriched in output (green rectangle) relative to input are typically considered selection hits. D) Scatter plot of normalized output versus input read counts against control (streptavidin). E) Flow chart of panning campaign against NS3a* with baseline. F) Procedure of final round selection with a spiked-in non-binding control library (gray). G) Library composition after selection between target and control. After selection on the target, the baseline library is nearly absent (<1%), indicating strong selective enrichment on the target. H-I) Scatter plot of output vs input reads after normalization. Gray dots represent baseline peptides. J-K) Bar plots of peptide enrichment after binning by input abundance. The y-axis shows log (FC+0.001), where FC is fold change (output/input), to allow visualization of peptides with low or zero abundance in output. L-M) histogram of peptide enrichment originated from input reads bin 10-32 (L) and 11-38 (M).

We then repeated the same selection with a baseline control spiked prior to round 3 selection (**Fig. 1E-F**). The baseline was a random sample of peptides with isosteric population of disulfides and orthogonal silent barcode. Baseline and R3 input were mixed in a 70:30 ratio prior to round 3 of selection (**Fig. 1F**). Analysis of round 3 after on target selection showed that the baseline library represented only 0.04% of total reads (**Fig. 1G**) indicating a 70/0.04 = 1750-fold depletion when compared to peptides pre-selected to bind to NS3a*. In the control selection, baseline library represented 50% of the reads, and 70/50 = 1.4-fold depletion (**Fig. 1G**). The scatter plot of the NGS reads highlighted substantial difference between the “target” and “control” selections. In “control” selection baseline coincided with on-diagonal population (**Fig. 1I**). In selection on NS3a*, baseline peptide sequences were separated from the “diagonal” population by several orders of magnitude (**Fig. 1H**). Red rectangle highlights both R3 and baseline peptides that have the same copy number in the input (∼100 parts per million); after selection on NS3a*, few baseline peptides have CPM=1-3 and most have CPM<0; whereas for all R3 peptides CPM ranges from 20 to 1000. A selection with baseline (**Fig. 1H**) unveiled an observation missed in the selection without the baseline (**Fig. 1C**): an average peptide in R3 is a significantly stronger binder to NS3a* than a random population of peptides. The fundamental lesson is that peptides that do not exhibit enrichment in late rounds of multi-round selection cannot be equated to non-binders. This observation is a classical red queen effect: the peptides need to interact significantly with the NS3a* simply to retain their constant position in the library. Baseline peptides that do not bind to NS3a* are falling behind or are removed from the selection (CPM<0).

To quantify whether enrichment is associated with specific sequence motifs, we employed Bioconductor-DE analysis (14, 46) to calculate the fold change (FC) from output and input reads for each peptide. Reads were binned based on their input abundance, and FC was plotted across bins. We separated peptides with DMT (yellow bars) from those that contain no DMT motif (blue bars) from “baseline” (grey bars). During selection on NS3a*, both DMT-containing peptides and non-DMT sequences were separated from the baseline. Although DMT-containing peptides exhibited higher median FC values in each bin (**Fig. 1J and L**), there existed a clear population of non-DMT peptides that enriched several orders of magnitude above the baseline. These observations show that focusing analysis to recognizable peptide motifs (47) may be counterproductive. Indeed, we have previously demonstrated that non-DMT sequences from NS3a* selection have confirmed binding to NS3a* (45). In the control selection, FC of DMT-containing peptides and non-DMT sequences was indistinguishable from the baseline population, indicating lack of binding to SMB (**Fig. 1K and M**).

To demonstrate the use of baseline in evaluation of robustness of the selection, we compared NS3a* selection from disulfide libraries to the same selection that employed libraries chemically modified by metabromoxylene (MBX). Spiking a baseline population at the third round of selection uncovered an intriguing observation that MBX-modified library exhibited an inferior selection outcome. In the R3 of the selection, the enrichment factors of >90% of the population of MBX-modified macrocycles were indistinguishable from enrichment of random baseline population (**Fig. S1D**). We compared this data to the enrichment factors of peptides of the same abundances the R3 of selection on NS3a* that employed unmodified disulfide macrocycle: 99% of the powered clearly separated from the baseline population (**Fig. S1C**). The most abundantly present peptides at R3 in both selections contained DMT-motifs (**Fig. S1C-D**), hence, motif-based heuristic assessment of this discovery campaigns can deemed both of them as “success”. Nevertheless, calibration of these selections against the same baseline uncovered significant decrease in the enrichment of the MBX-modified library because only a minor subset of sequences in R3 exhibited an enrichment above the “baseline response”. This suboptimal performance might be the result of the known ability of certain cysteine-reactive crosslinkers such as MBX to decrease the infectivity of phage (48); and it is possible that compromised infectivity of phage limits the efficiency of multi-round selection.

### Forging baseline in selection by mixing selections from unrelated targets

A binned analysis in a previous section (**Fig. 1J-K**) illustrates that a robust baseline should not only contain a random population of peptides, but also exhibit peptide abundances (measured as counts per million (CPM)) that resemble those of the preselected populations. Such CPM-matched random baseline is not always readily available. A sample of the naïve library does not form robust baseline because all peptides in such population have low CPM. We hypothesized that in selection on unrelated targets T1 and T2, selected population S1 can serve as a baseline and selection for target T2 and vice versa. A peptide population selected on unrelated targets is likely to contain peptides with high and low CPM and, in principle, it serves as a functional baseline. Blending of unrelated selections is counterintuitive: why contaminate the library with unrelated binders after it has been carefully selected to enrich only useful sequences? Nevertheless, the “in silico mixing” of hits from unrelated campaigns is an acceptable practice: for example, a recent preprint from Dickinson and co-workers trained contrastive learning models by combining data from binding campaigns on protein P1 (“binder” population) and data from a separate binding campaigns on unrelated protein P2: peptides enriched in P2 and not P1 campaign we denoted as “non-binder” population (35).

To test the benefits of physical mixing of unrelated libraries, we conducted a separate phage displayed campaign starting from SX₂CX₈CX₂ library to identify peptides that bind polysaccharide chitin (**Fig. 2A**). This selection employed chitin beads and traditional acid elution. In short, the selection was successful because R3 library clearly separated from the baseline population (**Fig. 2B**). A dominant HPV motif emerged among the top enriched peptides (**Fig. S2A**) and a synthetic biotinylated chitin-binding peptide isolated from the campaign bound to chitin magnetic beads as measured by fluorescence (**Fig, S2B-C**), and chitin-like glycans but not to unrelated glycans in glycan array (**Fig. S3**). Given the distinct molecular characteristics of chitin (a polysaccharide) and NS3a* (a protein), we postulated that libraries selected against these targets may thus serve as mutual baselines. To test the cross-target baseline concept, we blended SX₂CX₈CX₂ library from R3 input of chitin selection and the SX₃CX₉C library from R3 input of NS3a* selection and further spiked a *bona fide* random baseline used above. The combined library was used as input for round 3 selections against either chitin or NS3a* (**Fig. 2B**). In selections against NS3a*, the average fold change of peptides from the chitin-selected library closely matched that of random baseline across all input bins (**Fig. 2B**), indicating that chitin-binding peptides do not exhibit specific binding to NS3a* and functioned effectively as a baseline. In the reciprocal selection against chitin, the SX₃CX₉C library (pre-selected for NS3a*) was indistinguishable from bona fide baseline (**Fig. 2C**) whereas chitin-selected population contained populations of peptides that were clearly separated from both baseline populations. These results justify the blending of libraries derived from unrelated selection campaigns as practical implementation of baselines. Encouraged by these results, we push the cross-target baseline concept to campaigns with 22 protein targets.

**Figure 2.**
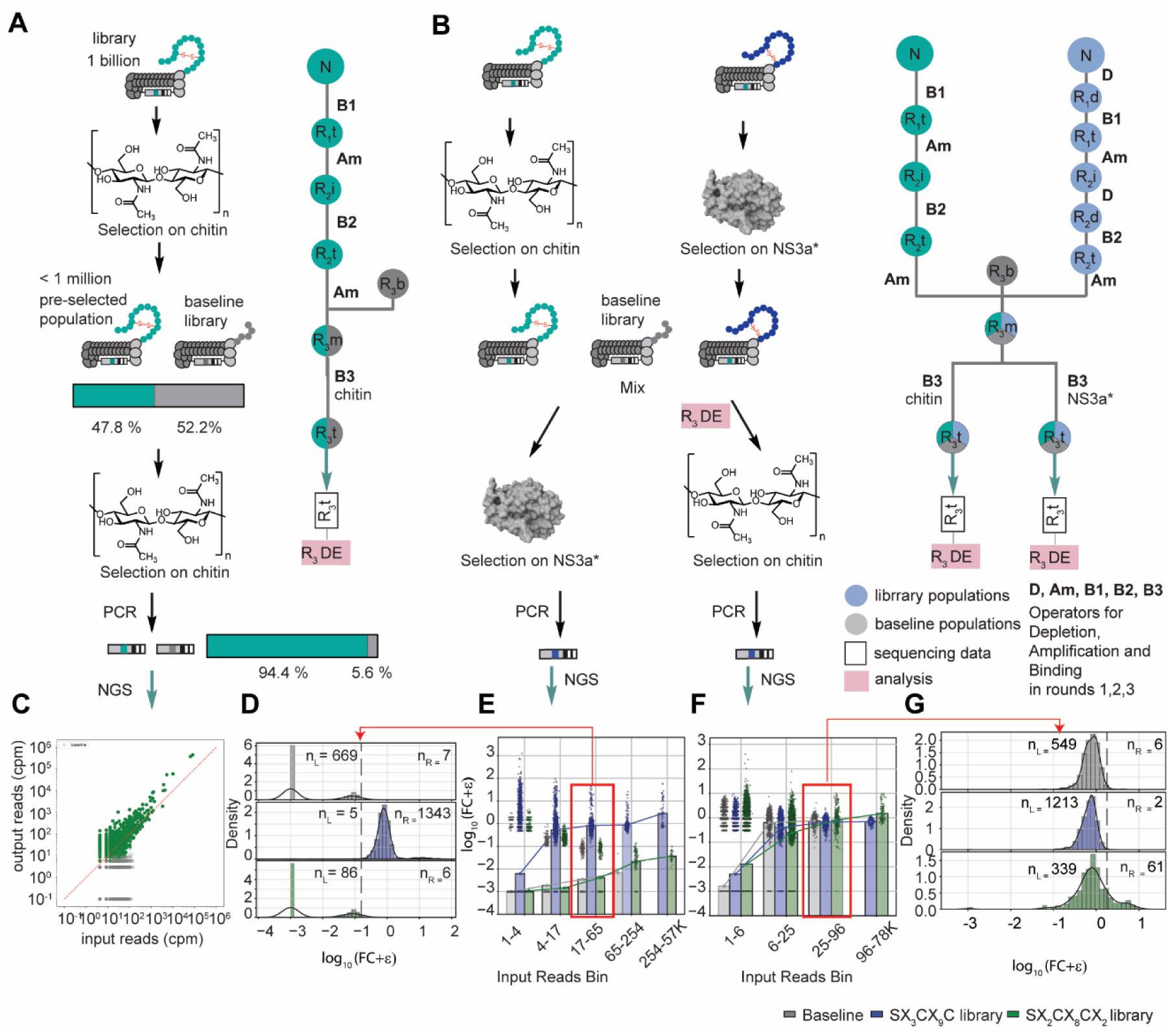
Using library selected for a different target as baseline. A) Procedure of selection on chitin (a polysaccharide) with baseline B) Procedure of selection on structurally different targets: NS3a* (a hepatitis C viral protease) and chitin. In each case, in round 3, the target-specific library was mixed with two controls: a non-binding baseline and a library previously selected on a different target. Selections were then performed separately on NS3a* and chitin. C) Scatter plot of normalized output versus input read counts against chitin. D) Histogram of peptide enrichment originated from input reads bin 17-65. E-F) Bar plots showing enrichment after binding by input abundance. In the NS3a* selection (E), the NS3a*-enriched library displayed strong divergence from both the non-binding baseline and the chitin-selected library, indicating target-specific enrichment. In contrast, during selection on chitin (F), the three libraries (chitin-selected, NS3a*-selected, and baseline) showed similar behavior, suggesting minimal selective enrichment and poor discrimination between peptide populations. G) Histogram of peptide enrichment originated from input reads bin 25-96. Note that in the highest frequency bin, the peptide from *bona-fide* populations were missing but NS3A*-selected peptides offered a convenient baseline response.

### Use baseline in semi-automated analysis of parallel selection campaigns

Presence of constant baseline in the molecular discovery procedure makes it possible to standardize the analysis of the data emanating from multiple selections. As an illustration, we performed 66 parallel selections using 22 protein targets and 3 phage-displayed libraries (**Fig. 3A-D**). Each protein was expressed by a commercial supplier as a fusion with a His-Avi tag for oriented immobilization on streptavidin or Ni-NTA. Most cell-surface proteins in this campaign are targets of interest for development of peptide-based radiopharmaceuticals (49). In the first round of selection (R_1_), targets were immobilized on the surface of streptavidin-coated 96-well plate and incubated with the phage-displayed libraries (**Fig. 3A**). After wash and elution and amplification, the libraries of different architecture selected on the same target were pooled and used as an input for the second round of selection (R_2_ **Fig. 3A-B**). The R_2_ employed protein immobilized on Ni-NTA beads (**Fig. 3B**) and an automated bead-washing, elution, and amplification workflow to produce R_3_ input for each target. Instead of using them individually, we combined R_3_ inputs from multiple selection. These combined inputs were employed in the third round of selection (**Fig. 3B-C**). The NGS data from R_3_ input and output from panning on the target and control (streptavidin beads without target) were processed using DE algorithm that identified a population of peptide enriched above the baseline population in the target but not control selection (**Fig. 3D**). As we observed in previous sections, mixing R_3_ inputs emanating from selection campaigns on distinct targets forms an effective baseline; however, a prior experiment carefully delineated library architectures and did not exhaustively test the limits of such mixing. In this selection we mixed R_3_ inputs from 22 targets in batches of N=4-12 campaigns to test whether such mixing can yield a productive baseline (**Fig. 3E**). Automation of the DE analysis was one of the benefits of mixing the inputs from as many targets as possible: in the selection from the mixed population the DE analysis recycles the data from input library and control selections. We performed validation of 16000 peptides in two tiers using focused library (Tier 1) and validation of binding properties of 27 synthetic peptides was done using surface plasmon resonance and ELISA (Tier 2).

**Figure 3.**
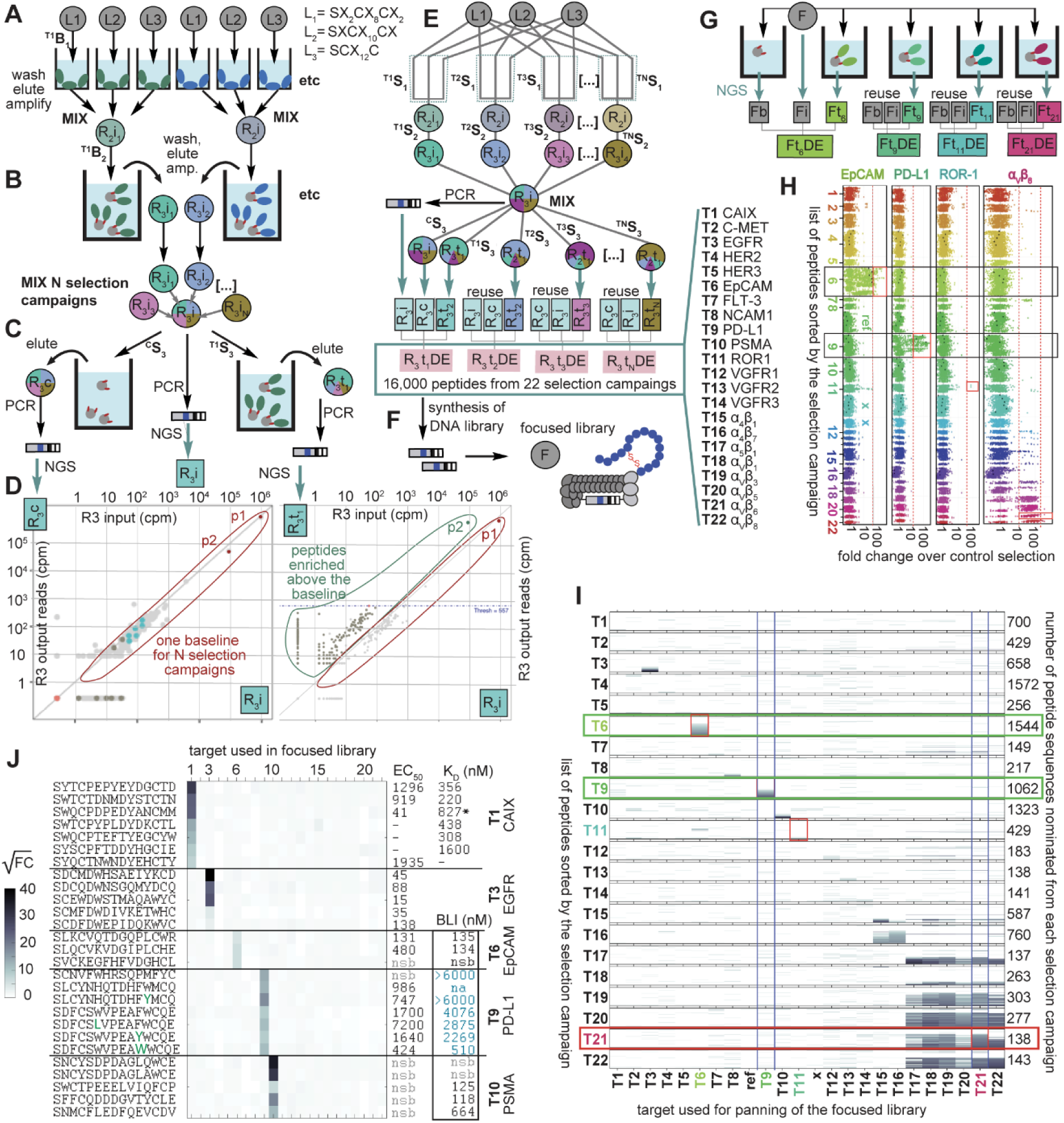
Application of baseline-based detection in parallel panning against multiple targets. A) Example of workflow showing selection of three different libraries against 2 targets using 3 libraries. B) Selection for the same target originating from different libraries are mixed at round 2. C) At Round 3, multiple selection campaigns are mixed and used as input for Round 3 for multiple targets and control selections. D) example of differential enrichment (DE) analysis that identifies baseline sequences and sequences enriched above the baseline in panning against a specific target. In this workflow, multiple selection campaigns share the same baseline. E) Mixing of multiple campaigns and universal baseline makes it possible to streamline analysis of 3 phage libraries against 22 targets (66 independent selections progressively mixed together). F) The same DE-analysis algorithm can be iteratively applied to all selection campaigns to nominate the hit compounds (16,000 total) which were validated by testing them all in parallel in a focused library. G) panning of the same focused library against the targets, controls, followed by DE of NGS data identified enrichment profile for the focused library in each selection presented as a Manhattan plot (H) or as a heat-map (I). For example in selection against target 6 (EGFR) or T9 (PD-L1) enriches subset of peptides from 1544 peptides nominated from T6 or T9 selection campaign. In contrast, peptides integrin selection campaigns show considerable cross-reactivity.

In Tier 1 validation, we used 16,000 peptides identified by DE from 22 selections against targets T1-T22. They were converted to a DNA library encoding such peptides and cloned into a phage displayed focused library [F], as described in our previous report (46). Binding of the [F] to the original 22 targets (**Fig. 3G**) followed by NGS and DE-analysis identified peptides that exhibited a significant fold change (FC) enrichment when compared to the input and control selections (**Fig. 3E**). We observed three types of outcomes. For targets like EpCAM (**T6**) and PD-L1 (**T9**), selections can be marked as successful. A significant fraction of the peptides nominated to bind to these targets (i.e. [T6^O^] and [T9^O^]) indeed have been confirmed to bind to these and only these targets (**Fig. 3H**). From 1544 peptides from [T6^O^], those discovered in panning of T6 (EpCAM), 500 bind to EpCAM above the baseline, but not to PD-L1, ROR-1 or integrin avb6. Similarly, peptides from [T11^O^] population bind to PD-L1 but not to EpCAM, ROR-1 or integrin. In **Fig, 3I,** which is a heatmap summary of 22 Manhattan plots describing 22 focused library experiments, it is evident that peptides selected in T6 or T9 campaigns that bind to T6 or T9 do not bind to any other 21 targets. For ROR1, from 429 peptides nominated to bind to ROR-1 only 2 peptides exhibited binding to ROR-1 above the “baseline” (**Fig. 3H**). Finally for integrin targets, we observed an expected poly-specific binding. Binding of [F] to α_V_β_6_, identified binders not only from [T21^O^] but also from [T16^O^]-[T20^O^] and [T22^O^]. From 138 peptides in [T21^O^] population many peptides also bind to T17-T20 and T22 (red box, **Fig. 3I**). In general, selection for T17-T22 targets often identified peptides that bound to the cognate target and other targets in this cluster, but not to other proteins T1-T16. Selection for T16 (α_4_β_7_) identified peptides that cross-reacted with T15 (α_4_β_1_) but not T17-T22 integrins. Selection for T15 (α_4_β_1_) identified many peptides that cross-reacted with T18 (α_V_β_1_) and a few peptides cross-reacting with T16 (α_4_ β_7_), T19 (α_V_ β_3_), T21 (α_V_β_6_) T22 (α_V_β_8_).

In Tier 2 validation, we synthesized 27 peptides from 5 selection campaigns with C-terminal biotin and validated their interaction with the immobilized target using ELISA and dose-response titration of soluble biotinylated peptide to estimate EC_50_ of binding. We employed at least two other assays. When limited solubility biotinylated peptides or high non-specific binding (nsb) prevented ELISA measurements (e.g., PSMA or EpCAM, **Fig. 3J**), we employed bilayer interferometry (BLI) assay to estimate K_d_ using immobilized biotinylated peptide and soluble protein, For CAIX we re-synthesized some peptides without C-terminal biotin and re-tested them by surface plasmon resonance (SPR) with immobilized CAIX to yield kinetic K_d_ values (**Fig. 3J**). Overall, the Tier 2 validation confirmed that most peptides that exhibit significant FC values in focused library exhibit mid-nanomolar to single digit micromolar potency in the binding assays.

In the parallel selection against 22-target we did not employ any cross-target depletions. Emergence of poly-specific hits in such selections is anticipated outcome, and has been well-documented in phage display (50) and even in continuous selection procedures (51). A selection against 22 targets covered an array of possibilities from unrelated targets to weakly related HER1-HER3 (**T3**-**T5**), VEGFR1-3 (**T12**-**T14**), integrins with weak homology (**T15** vs. **T17**-**T22**) to close homologues (**T17**-**T22**) to extreme homologues such as **T18** and **T22** that differ in only few amino acids in the vicinity of the peptide binding site. We anticipated that by mixing selections from unrelated targets (**T1**: carbonic anhydrase 9 or CAIX), **T2** (c-Met), **T3** (EGFR or HER1), etc. the peptides selected to bind to c-Met and EGFR would form a “baseline” relative to CAIX because these three targets share no homology. It is unlikely for c-Met-binding peptide or EGFR-binding peptides to bind to CAIX. Indeed, these selections proved to yield binding peptides that confirmed binding to these and only these targets in focused library selection (**Fig. 3I**). In the selections against homologous integrin targets (**T17**-**T22**), selection of cross-reactive hits (**Fig. 3I**) was not surprising. For example, selection of peptides against α_4_β_7_ often exhibit cross-reactivity to α_4_β_1_. Selections that use integrins α_V_β_3_, α_V_β_5_, α_V_β_6_ by phage display have been documented (52–54). It is known that outcomes from selection on these integrins frequently give rise to peptides that bind to homologous integrins and selection of isoform-specific peptides requires intricate depletion against cross-reactive targets.(16, 55) Beyond cross-reactivity, the selection procedure successfully identified 60-80% of peptides that bound to the cognate integrin but not to T1-T16 targets outside integrin family. Cross-reactivity with other integrins is unlikely to be related to mixing of libraries or use of baseline in DE-analysis. Selection and validation of integrin-specific peptides that avoid binding to closely related homologues are beyond the scope of this publication.

We tested whether baseline-stratified FC values can offer decision enabling information for the optimization of the properties of sequences. We focused this questions on sequences discovered to bind to PD-L1 protein. The focused library F contained 12×18=216 sequences derived from peptide S<u>DF</u>C<u>SWVPEAFW</u>C<u>QE</u> in which every underlined position was substituted by 18 other amino acids (apart from Cys and the parent amino acids). The heatmap in **Fig. 4A** describes the FC enrichment of each derivative sequence in the panning experiment. We noted that the changes in certain position gave rise to a baseline-level FC (**Fig. 4A**). This loss of function indicating that a specific amino acid is indispensable for binding. In other positions several substitutions could be justified as “neutral changes” or improvements (**Fig. 4A**). The analysis of absolute of relative FC values (**Fig. S4**) suggested at least six positions for optimization; a separate deep-mutational scan (**Fig. S5**) suggested further modifications of the N and C termini of the S<u>DF</u>C<u>SWVPEAFW</u>C<u>QE.</u> These plausible positional changes (**Fig. 4B**), when applied individually or in combination to S<u>DF</u>C<u>SWVPEAFW</u>C<u>QE</u> gave rise to a series of sequences that exhibited a single digit nanomolar potency in cell binding and SPR assays (**Fig. 4C**) (**Fig. S6-S12**). We note that improvement over that 3-4 orders of magnitude in binding performance (**Fig. 4C**) was possible without engagement of non-canonical amino acids (ncAA). The only exception was the use of 4R-fluoroproline denoted as π in Fig. 4C. We noted that 4R-fluoroproline provided 2-3-fold benefit in binding whereas 4S-fluoro proline in the same position significantly diminished the binding (not shown). These improvements likely result from the stereo electronic role of fluorine in attenuation of the equilibria between exo/endo ring pucker and trans/cis conformational of the amide bond in proline (56). The properties of the identified sequences with single digit nanomolar performance in cell binding assays were further improved by ncAA mutagenesis but the discussion of these improvements is beyond the scope of this manuscript and they will be presented elsewhere.

**Figure 4.**
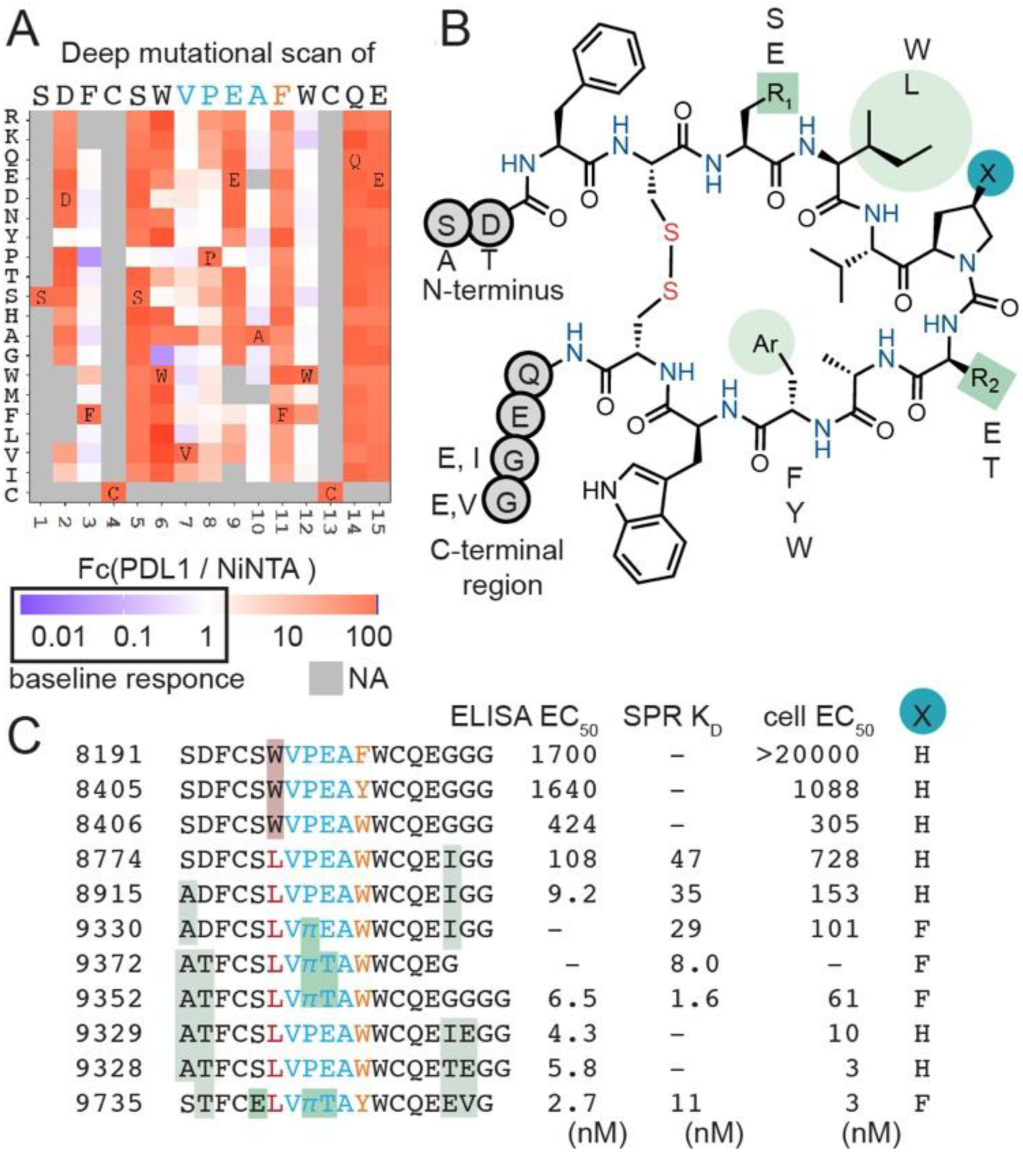
Application of baseline-based measurements of enrichment as decision-enabling information in optimization. A) Heatmap describing changes in binding performance of phage displayed peptides, as measured by FC, caused by systematic changes of every amino acids to every other proteogenic amino acid (aka “deep mutational scan”). B) Structure of the core region with key optimizable positions identified by the deep mutational scan. C) Synthetic peptides with individual chances of combinations of changes suggested by the deep mutational scan. Properties of the peptides have been assessed in three independent assays. ELISA assay employed a dose-response titration of soluble biotinylated peptide and PD-L1 immobilized on microtiter well place. SPR employed a surface immobilized PD-L1 and soluble peptides; cell-based assay employed a dose response titration of biotinylated peptide with PD-L1(+) CHO cells, where the binding of the peptides was probed by fluorescent streptavidin.

## Discussion

The advent of next generation sequencing (NGS) and introduction of NGS to phage display and other selection procedures around 2010’s encouraged many research groups to postulate that a “deep” analysis of *in vitro* selections by NGS holds a promise to identify hit sequences missed from *in vitro* selection procedures in traditional analyses based on Sanger sequencing of isolated clonal populations. Delivery on this promise requires a unified framework for the mathematical analysis of the NGS data emanating from the multi-round *in vitro* selection. Our observation is that such unified framework is still lacking. Some semi-quantitative approaches combine analysis of NGS frequency with analysis of peptide motifs (45, 57); algorithms that search for motifs in NGS data in a supervised or unsupervised fashion have been reported (58, 59); most algorithms have not been adopted across the community. The other analysis simply states that the most abundant peptides in the NGS should be prioritized for validation. Other arguments prioritize sequences that change their enrichment in two adjacent rounds (60). The latter could be tainted by the amplification bias in multi-round selections (14). Bioconductor DE pipeline uniformly accepted in RNASeq community offers unbiased statistics-based analysis of sequences significantly enriched between the test and a plurality of control populations. This pipeline can automatically process hundreds of datasets in parallel. Our lab repurposed Bioconductor DE pipeline to analyze amplification bias in phage libraries and identify enrichment in phage display (61) in DNA-barcoded population of peptide derivatives (44), glycans and other molecules that fail in clustering approaches (62–65). Unfortunately, a minor yet fundamental difference exists between RNASeq and *in vitro* selection datasets. All RNASeq datasets contain a *bona fide* baseline formed by non-responder transcripts. Such baseline is often absent from *in vitro* selection procedures. In our lab, we colloquially refer to it as a “swimmer paradox”: DE analysis can readily delineate fast and slow swimmers, but DE analysis fails in analysis of non-swimmers and DE fails in the analysis of the population of Olympic swimmers if they finish the lap with nearly matched time. Without an absolute reference (e.g., chronometry) DE cannot distinguish a uniform population of Olympic swimmers from an equally uniform population of individuals whose swimming ability is zero. Similarly, in a population of strongly binding peptides, DE-analysis fails if all peptides exhibit similar strength of binding. Failed DE in local analysis cannot distinguish between specific and non-specific outcomes and we documented failed DE-Analysis in successful selection procedures (46). Introducing a baseline into all *in vitro* selections makes DE-analysis robust because binding performance of all sequences is stratified against an absolute non-binding reference. Mixing libraries from two selection procedures breaks the swimmer paradox because it places Olympic swimmers and non-swimmers in the same pool. Our manuscript demonstrates that introduction of baseline enables robust identification of true binders as those that outperform randomly distributed or irrelevant peptide sequences. Baseline allows robust stratification of the selection procedures and automated analysis of large number of selection procedures in parallel.

## Materials and Methods

### General biochemistry information

The libraries were constructed using TriNuc codons that only contains 19 specific codons excluding cysteine (66). The procedures have been adopted and modified as previously described in publications that produced the M13-displayed SXCXXXC library (61), M13-SDB vector (67), SXCX_4_C and SXCX_5_C libraries (43). In short, to produce SX_3_CX_9_C library, the vector SB4 QFT*LHQ was digested with *KpnI* HF (NEB Cat# R3142S) and *EagI* HF (NEB Cat# R3505S). A primer/template pair consisting of annealed primer 5′-AT GGC GCC CGG CCG AAC CTC CAC C-3′ and template 5′-CC CGG GTA CCT TTC TAT TCT CAC TCT TCT **XXX** TGT **XXXXXXXXX** TGT GGT GGA GGT TCG GCC GGG CGC TTG ATT-3′ with ‘XXXXX’ representing trinucleotide cassette (TriNuc) that contains a mixture of 19 codons that encode 19 natural amino acids except cysteine (oligos containing the TriNuc mixtures was synthesized by Genscript). The primer/template was then extended using Klenow DNA polymerase (NEB) according to the manufacturer’s instructions. The insert fragment was then digested with *KpnI* HF and *EagI* HF, gel purified and ligated into the cut vector. The ligation products were then transformed into electrocompetent E. coli cells, and the transformants were grown overnight on *E. coli* TG1 to allow for phage production. Phage cultures were then centrifuged to remove cells and debris, and then the phage was precipitated by PEG precipitation (5% PEG 0.5 M NaCl).

In our previous reports, we characterize diversity of small libraries fully by NGS (61, 68). The true diversity of 10^9^ scale library is difficult to estimate because it would require a prohibitively expensive NGS of >10^10^ reads (10× coverage of diversity), e.g. requiring 400 runs with MiniSeq High Output Reagent Kit with 25 million reads each, costing 400×$3000 = $1,200,000.

Based on our previous reports, diversity can be inferred from NGS analysis of the sample of naïve libraries, for example, SX_3_CX_9_C library covers 2,309,529 reads and detects 2,268,469 unique sequences.

The SX_3_CX_9_C, SX_2_CX_8_CX_2_ and SDB 17/SDB10 GS23 libraries were obtained from 48HD. The sequencing files for naïve libraries were uploaded to https://48hd.cloud/ and the links are attached below.

SX_3_CX_9_C: https://48hd.cloud/file/11581

SX_2_CX_8_CX_2_: https://48hd.cloud/file/11471

GS23: https://48hd.cloud/file/18801

### Panning procedures on NS3a*

Panning procedures on NS3a* was same as described in previous publication.(45)

### Panning procedure on chitin

Panning procedures were performed in 3 rounds. In all rounds, chitin magnetic beads (New England Biolabs, #E8036S) were used as the target. Blocking, binding and washing steps were done in KingFisher™ Duo Prime Purification System (Thermo Scientific™, #5400110). Chitin magnetic beads were vortexed for 2 seconds and the solution was attached to the magnet to remove the storage buffer. The beads were washed three times with binding buffer (20 mM Tris HCl, 500 mM NaCl, 1mM EDTA disodium, pH 8.0) and resuspended in 100 uL binding buffer.

For round 1, the beads suspension and other reagents were added to a 96 Deepwell Plate (Thermo Fisher, #95040450) as follows:

Row A: Chitin magnetic beads (New England Biolabs, #E8036S) (0.1 mL/well)
Row B: reserved for 12-tip Deepwell magnetic comb (Thermo Fisher, #97003500)
Row C: Binding Buffer (20 mM Tris HCl, 500 mM NaCl, 1mM EDTA disodium, pH 8.0)
Row D: Blocking Buffer (1 mL, 2% BSA (w/v) in Binding buffer)
Row E: Solution of S2C8C2 phage library (100 μL, 10^12^ PFU/mL)
Row F: Wash Buffer (1 mL, 0.1 % Tween in Binding buffer)
Row G: Wash Buffer (1 mL, 0.1 % Tween in Binding buffer)
Row H: Binding Buffer (1 mL, 0.1 % Tween in Binding buffer)

Following steps were performed using a KingFisherTM Duo Prime Purification System with a magnetic comb to transfer the beads. The program is as follows:

a) Collect comb from row B
b) Collect beads from row A on comb
c) Wash beads in row C – 30 s
d) Block in row D – 1 h
e) Phage binding in row E – 1.5 h
f) Wash beads in row F – 1 min
g) Wash beads in row G – 1 min
h) Wash beads in row H – 1 min.

At the end of the program, the protein coated beads with phage bound were in wells in the last row. The content of each well from row 8 was transferred to individual Eppendorf^TM^ tube and the tubes were placed in Dynabead^TM^ MPC-S for 30 seconds to capture the beads. The supernatant was discarded, the beads were resuspended in elution buffer (200 μL, 200 mM glycine pH 2.2) and rotated on Thermo Scientific Labquake 360 rotator (cat# C400220Q) for 9 min and then 30 uL of neutralization buffer (1M Tris HCl pH 9.1) was added immediately. Eluted phage bound was taken for further processing. 200 uL of eluted phages were used for phage amplification (See *Amplification* protocol).

For round 2, the same protocol was followed, except that we used 30 uL of chitin magnetic beads instead of 50 uL and three washes were performed in 1 mL, 0.1 % Tween in Binding buffer instead of two washes. For round 3, we included the baseline library, used 30 uL of chitin magnetic beads and performed 6 washes in 1 mL, 0.1 % Tween in Binding buffer

### Chemically modification of phage libraries

Macrocyclization with MBX: a solution of SX3CX9C phage library (10 µL, ∼10^13^ PFU/mL PBS in 50% glycerol, pH 7.4, amplified output of Round 2) was first diluted with 62 µL of water in a 1.7 mL Eppendorf^TM^ tube and 10 µL of 1M Tris-base buffer (pH 8.6) was added to adjust the pH of the reaction mixture to 8.6 (checked by universal pH paper). TCEP (2 µL, 100 mM in water) was added to the reaction, mixed by vortexing and incubated at room temperature for 30 min. The reaction was loaded on an equilibrated Zeba Spin column (Thermo Fisher cat# 89883) to remove excess amount of TCEP. 10 µL of 1M Tris-base buffer (pH 8.6) was added, and the macrocyclization reaction was initiated by the addition of MBX linker (1 µL, 10 mM in DMF). The reaction mixture was mixed and incubated at room temperature for 20 min. After 20 min, the reaction mixture was loaded on an equilibrated Zeba Spin column and eluted by centrifugation in 2000 xG. The macrocyclized phage library was obtained as a cleared colorless solution.

### Phage amplification

70 μL of eluted phages were mixed with 250 μL of E2773 bacterial culture and then added to 25 mL of LB media for amplification at 37 °C for 4.5 h. After amplification was done, the culture was transferred to 25 mL Falcon tube and centrifuged for 10 mins at 6,000 × g at 4 °C. Supernatant was transferred to a new 25 mL Falcon tube and 1/5 volume of PEG/NaCl (∼5 mL) was added. Phage was incubated 4 °C overnight. Next day, phages were centrifuged at 14,000 × g for 30 mins at 4 °C and the pellet resuspended in 1 mL PBS. The suspension was transferred to 1.7 mL microcentrifuge tube and centrifuged at max speed for 5 mins at 4 °C. The supernatant was transferred to a new 1.7 mL microcentrifuge tube and 1/5 volume to PEG/NaCl (∼200 µL) was added and incubated for 1 h on ice. Suspension was centrifuged at maximum speed for 30 mins at 4 °C and the the pellet was resuspended in 500 µL of PBS, and stored at 4 °C.

### PCR of phage

Two-step semi-nested PCR was used to improve the sensibility. The method was adapted from our previous developed protocol.(69, 70) DNA template (phage solution) was amplified in Phusion® HF buffer with Phusion® High-Fidelity DNA Polymerase (NEB, #M0530S).

1^st^ step:

A typical 50 μL reaction mixture contained:

1. 5x Phusion buffer 10 μL
2. 10 mM dNTPs 1 μL
3. DMSO 2.5 μL
4. Phusion® Polymeras 0.5 μL
5. Forward primer (TTTTGGAGATTTTCAACGTG, 10 μM) 1 μL
6. Reverse primer (CCCTCATAGTTAGCGTAACG, 10 μM) 1 μL
7. DNA Template solution 10 μL
8. Nuclease free water 24 μL

Cycling was performed using the following thermocycler settings:

a) 98 °C 3 min
b) 98 °C for 10 s
c) 50 °C 20 s
d) 72 °C 20 s
e) repeat b)-d) for 30 cycles
f) 12 °C 1 min
g) 4 °C hold

2^st^ step:

A typical 50 μL reaction mixture contained:

1. 5x Phusion buffer
2. 10 mM dNTPs
3. 10 μL 1 μL 2.5 μL
4. Phusion® Polymeras 0.5 μL
5. Forward primer (10 μM) 1 μL
6. Reverse primer (10 μM) 1 μL
7. PCR product from 1st step 2 μL
8. Nuclease free water 32 μL

Cycling was performed using the following thermocycler settings:

a) 98 °C 3 min
b) 98 °C for 10 s
c) 50 °C 20 s
d) 72 °C 20 s
e) repeat b)-d) for 20 cycles
f) 12 °C 1 min
g) 4 °C hold

### General data processing methods

Core scripts are available as part of the Supplementary_data.zip and https://github.com/derdalab/KYL. Data storage cloud http://48hd.cloud/ was implemented in Linux-Apache-MySQL-Python (LAMP) architecture and details of this implementation are beyond the scope of this report. We used Bioconductor EdgeR DE analysis with modeling of the observed counts using a negative binomial model, Benjamini–Hochberg (BH) adjustment to control the FDR at α = 0.05 and normalization of data across multiple samples using the TMM normalization. Prior to DE-analysis, “test” and “control” datasets were retrieved from the https://48hd.cloud/ server as tables of peptides, DNA, and raw sequencing counts. Tables of raw sequencing counts and DA values presented in this manuscript are available in Dataset S1.

### Processing of Illumina data

The Gzip compressed FASTQ files were downloaded from BaseSpace™ Sequence Hub. The files were converted into tables of DNA sequences and their counts per experiment. Briefly, FASTQ files were parsed based on unique multiplexing barcodes within the reads discarding any reads that contained a low-quality score. Mapping the forward (F) and reverse (R) barcoding regions allowing no more than one base substitution each and F-R read alignment allowing no mismatches between F and R reads yielded DNA sequences located between the priming regions. The files with DNA reads, raw counts, and mapped peptide modifications were uploaded to http://48hd.cloud/ server. Each experiment has a unique alphanumeric name (e.g., 20230515-1666OOooJUA-KJ) and unique static URL: for example https://48hd.cloud/file/14363)

URL links to sequencing data can be found in Supplementary Tables S1-S6. Sequencing files of the reshaped libraries were further processed by python to sort mixed libraries, combine repeated peptide sequences, remove stuffer and other peptides sequences that are not related to the libraries of use. The scripts and the processed sequencing files are available in Dataset S2.

## Acknowledgments

The authors acknowledge funding from Natural Sciences and Engineering Council of Canada (RGPIN-2022–04484 to R.D.), Natural Sciences and Engineering Council of Canada Accelerator Supporting (to R.D.), Canadian Institutes of Health Research (CIHR FRN: 168961 to R.D.) and Mitacs Elevate Fellowship (to G.M.L.). Infrastructure support was provided by the Canada Foundation for Innovation New Leader Opportunity (to R.D.) and National Institute of General Medical Sciences of the National Institutes of Health (R01GM145011 and F31GM155953 to F.B.M).

## Author Contributions

K.Y., V.A. and R.D. designed research; K.Y., G.M.L., T.B., Z.O., C.K., M.L.M. and K.M.performed research; F.B.M., and W.K., provided critical reagents; K.Y., G.M.L., G.B., and R.D. analyzed data; K.Y., G.M.L and R.D. wrote the paper.

