## Supporting Information for "Universal Baseline for *in vitro* Selection of Genetically Encoded Libraries"

[a.] Department of Chemistry, University of Alberta, Edmonton, AB T6G 2G2, Canada
[b.] 48Hour Discovery Inc., Edmonton, AB, T6G 2M9, Canada
[c.] Zealand Pharma A/S, Søborg, Denmark
[d.] Graduate Program in Biological Physics, Structure and Design, University of Washington, Seattle, Washington 98195, United States

* Ratmir Derda

**This PDF file includes:**

Supporting text
Figures S1 to S12
Table S1 to S5
SI References

**Other supporting materials for this manuscript include the following:**

Dataset S1
Dataset S2

### Supporting Information Text

#### Abbreviations

| | |
|---|---|
| BSA | Bovine Serum Albumin |
| dNTP | Deoxyucleoside Triphosphate |
| DNA | Deoxyribonucleic acid |
| DMF | Dimethylformamide |
| ESI | Electrospray Ionization |
| LB | Lysogeny Broth |
| LiGA | Liquid Glycan Array |
| LCMS | Liquid chromatography Mass Spectrometry |
| MALDI | Matrix-assisted Laser Desorption/Ionization |
| MOPS | 3-(N-morpholino)propanesulfonic acid |
| MeCN | Acetonitrile |
| NGS | Next-generation sequencing |
| PBS | Phosphate buffered saline |
| PCR | Polymerase chain reaction |
| PFU | Plaque forming units |
| TCEP | Tris(2-carboxyethyl)phosphine) |
| TOF | Time of Flight |
| TRIS | Tris(hydroxymethyl)aminomethane |

### Supporting Methods

*Protein expression*
Plasmid construction, bacterial expression, protein expression was same as described in previous publication.(1)

*Binding of LiGA to peptide immobilized on plate*
Liophilized chitin-binding peptide was dissolved in PBS buffer at a final concentration of 5 µg/mL and 100 µL (0.5 µg) was added to each well of a 96-well Pierce™ Streptavidin Coated Plates (Thermo Scientific™, #15121). The plate was covered with sealing tape (Thermo Scientific™, #15036) and incubated overnight at 4 °C. The following day, 100 µL 10 mM solution (pH 8.0) was added to the wells and incubated at rt for 1 h. The wells were then washed 3x by adding washing buffer (200 µL, 0.1% Tween-20 in HEPES) in the wells and discarding the solution by inverting the plate on top of a paper towel. After blocking with biotin and washing, 100 µL of LiGA ($10^9$ PFU/mL in HEPES) was added to the wells. The solution was incubated for 1h at rt and discarded by inverting the plate. The wells were washed 2x with washing buffer and 1x with HEPES (200 µL). To elute bound phage, 50 µL of HCl (pH 2.0) was added to the well, incubated for 9 min at rt, and the content of each well was transferred to an Eppendorf tube containing 25 µL of 5x Phusion HF buffer (NEB, #M0530S). The neutralized solution was used for titering and as DNA template for PCR and Illumina sequencing. Binding of clonal phages (GlcNAc and blank phages) was performed in similar fashion and used for titering.

*Multi-Round Protein Panning*
Phage libraries were diluted in blocking buffer (CM-HBS + 1% BSA + 0.1mM biotin if using streptavidin support) at $10^{12}$ PFU/ml. Magnetic beads used for depletion were washed 3 times with 500 µL PBS and the naïve libraries were added onto the washed beads, and rotated at room temperature for 1 h. Depending on the target protein of the experiment, different beads were used. 1 µg protein (1 replicate per library) was added to pre-coated streptavidin wells in a total volume of 100 µL with 1X PBS, and shaken for 1 h at room temperature. The plate was washed 1 time with 300 µL 1X PBS and 100 µL blocking solution was added to wells and shaken for 1 h at room temperature. The supernatant was removed and 100 µL of round 1 input was added and incubated at 4 °C overnight.

On the next day, the wells were washed 3 times with 300 µL wash buffer (CM-HBS + 0.1% Tween-20). 200 uL of elution buffer (0.2M glycine-HCl, 0.1% BSA, pH 2.2) was added to the wells and incubated with gentle shaking for 9 min at room temperature. The elution buffer solution was then transferred to a tube containing 30 µL of neutralization buffer (**Round 1 Output**). 200 µL of this solution was transferred to to 1.5 mL ER2738 bacterial culture for phage amplification and incubated in shaker at 37 °C for 3 h at 200 rpm. The cell culture was centrifuged at 4000 rpm at 4 °C for 15 min and 1/6 volume of 6X PEG/NaCl (300 µL) was added to the supernatant. The sample was incubated on ice and centrifuged at 5,000 g for 5 min. Phage pellet was resuspended in 500 µL 1X PBS (**Round 1 Amplification**), diluted in blocking buffer to $10^{11}$ PFU/ml, and used for depletion on beads as described earlier (**Round 2 Input**).

Target appropriate beads were washed 3 times with 500 µL 1X PBS. 1 µg protein was added to washed beads in a total volume of 100 µL with 1X PBS and rotated at 4 °C overnight or 1 h at room temperature. The beads suspension and other reagents were then added to a 96 Deepwell Plate (Thermo Fisher, #95040450) as follows:

Row A: Bead solution (1 mL/well)
Row B: reserved for 12-tip Deepwell magnetic comb (Thermo Fisher, #97003500)
Row C: CM-HBS Buffer
Row D: Blocking Buffer
Row E: Round 2 Input (100 µL, $10^{11}$ PFU/mL)
Row F: Wash Buffer (1 mL)
Row G: Wash Buffer (1 mL)
Row H: Wash Buffer (1 mL)

After the program was finished, sample with beads was transferred from last wash buffer Row H to 1.7 mL tubes on magnetic rack to separate solution from beads. 200 µL of wash buffer was added to the beads and rotated for 9 min at room temperature. The elution buffer solution was transferred to a tube containing 30 µL of neutralization buffer (**Round 2 Output**). 200 µL of the Round 2 output were added back to beads and then added to 1.5 mL ER2738 bacterial culture for phage amplification following previous protocol for round 1 (**Round 2 Amplification**). The phage library was diluted in blocking buffer to $10^{10}$ PFU/ml and depleted as previously described (**Round 3 Input**).

Target appropriate beads were prepared by washing 3 times with 500 µL 1X PBS. 1 µg of protein and control protein, if applicable, were added to washed beads in a total volume of 100 µL with 1X PBS and rotated at 4 °C overnight. The beads suspension and other reagents were added to a 96 Deepwell Plate (Thermo Fisher, #95040450) as follows:

Plate 1:
Row A: Bead solution (1 mL/well)
Row B: reserved for 12-tip Deepwell magnetic comb (Thermo Fisher, #97003500)
Row C: CM-HBS Buffer
Row D: Blocking Buffer
Row E: Round 3 Input (100 µL, $10^{10}$ PFU/mL)
Row F: Wash Buffer (1 mL)
Row G: Wash Buffer (1 mL)
Row H: Wash Buffer (1 mL)

Plate 2:
Row A: Wash Buffer (1 mL)
Row B: Wash Buffer (1 mL)
Row C: Wash Buffer (1 mL)

After the program was finished, samples with beads were transferred from last wash buffer Row C, plate 2, to 1.7 mL tubes on magnetic rack to separate solution from beads. 30 µL of water was added to the beads and boiled at 90 °C for 10 min. The solution was then transferred to a new 1.7 mL tube (**Round 3 Output**).

*Focus Library Cloning*
PCR was used to produce double-stranded focused library from a DNA library oligos synthesized by GenScript. The DNA oligo was amplified as follows:

1. 10x Taq buffer 12 µL
2. 10 mM dNTPs 2 µL
3. Taq Polymerase 2 µL
4. GSX/IDT Oligo FW (10 µM) 3 µL
5. GSX/IDT Oligo RV (10 µM) 3 µL
6. Library Oligo 2 µL
7. $MgCl_2$ (25 mM) 8 µL
8. Nuclease free water 85 µL

Cycling was performed using the following thermocycler settings:
a) 95 °C 60 s
b) 95 °C for 20 s
c) 54 °C 20 s
d) 72 °C 15 s
e) repeat b)-d) for 35 cycles
f) 72 °C 30 s

The PCR reaction was purified using Macheney-Nagel PCR clean up kit using an adapted protocol. The phage vector was digested overnight with *KpnI* HF (NEB Cat# R3142S) and *EagI* HF (NEB Cat# R3505S). The insert PCR fragment was then digested with *KpnI* HF and *EagI* HF, purified using Macheney-Nagel PCR clean up kit and ligated into the cut vector. The ligation products were then transformed into electrocompetent *E. coli* 10G prlA4 cells, and the transformants were grown overnight on *E. coli* TG1 prlA4 cells to allow for phage production. Phage cultures were then centrifuged to remove cells and debris, and then the phage was precipitated by PEG precipitation using our standard phage purification protocol.

*ELISA*
ELISA measurements were performed on Pierce Nickel Coated Plate (Clear, 96-Well, 15442, Thermo Fisher) at room temperature. Plates were coated with his tag protein in 100 µL (0.5 µg/well) PBS, pH 7.4 for 1 hour at room temperature. Coated plates were washed three times with 300 µL of PBS and subsequently blocked with 200 µL SuperBlock (37515, Thermo Fisher) for 10 mins. Plates were washed again three time with PBS, + 0.05% Tween-20 (PBST). Then desired concentrations of serially diluted peptides solution (100 µL/well) in assay buffer (PBS, 0.05% Tween-20 & 0.5% BSA, pH 7.4) were added and incubated for 1 hour. For subsequent steps, ELISA plates were washed with PBST (300 µL, three times) in between the steps. Streptavidin-HRP (1:2000, STN-NH913, Acro Biosystem) was added to the plates and incubate for 45 minutes followed by TMB substrates, and 1 M phosphoric acid. Plates were promptly read for absorbance at 450 nM

on a microplate reader (Multiskan SkyHigh Spectrophotometer, Thermo Fisher). Data analysis and $EC_{50}$ values were calculated using GraphPad Prism (v.9.2).

*SPR*

Surface Plasmon Resonance (SPR) measurements were conducted using a Biacore 1K Instrument at 25 °C in PBS-P+ (0.2M phosphate buffer, 27 mM KCl, 1.37 M NaCl, 0.5% Surfactant P20, pH 7.4, Cytiva) with 0.05% DMSO. His tag protein was immobilized onto a Series S Sensor Chip NTA (BR100532, Cytiva) using the optimized concentration 5 µg/mL in assay buffer. Standard Biacore NTA immobilization condition was followed using NTA Reagent Kit (28995043, Cytiva). Following immobilization, peptides affinity measurements were made with the serially diluted solution in assay buffer using single cycle kinetics. The association and dissociation phases were monitored for 180 and 300 seconds respectively, with a flow rate of 30 µL/min. The chip was regenerated in assay buffer with 350 mM EDTA at flow rate 30 µL/min for 60 sec followed by 0.5 mM $NiCl_2$. In some measurement drift in signal was addressed using the amine coupling in NTA chip. Obtained data were analyzed with the Biacore Insight Evaluation Software and $K_D$ values estimated from 1:1 binding kinetics fit model.

*BLI*

The BLI procedure was performed as previously described, with minor adaptations (2).

*Cell binding*

Target cells expressing the protein of interest on the surface were collected and washed twice in CM-HBS buffer (50 mM HEPES, 150 mM NaCl, 1 mM $CaCl_2$, 1 mM $MgCl_2$) with 0.5% BSA. Washed cells were resuspended in ice cold Staining Buffer (SB, CM-HBS with 5 % FBS) to a concentration of $1x10^6$ cells/200µL and 200µL of cells were aliquoted into each tube (1 tube per data point). Peptides (biotinylated or fluorophore tag) solution was prepared to the desired starting point in SB from 10 mM DMSO stock, and then, serially diluted to lower concentration based on the expected binding affinity range. After labelling the tubes, supernatant was removed, and 200uL of the corresponding peptides concentration was added to each tube and incubated for 1 hour at 4 °C. During incubation time, cells were disturbed every 10-15 mins and then washed three times with 500 µL SB. For biotinylated peptides, cells were further incubated with Streptavidin-Alexa Fluor 488 Conjugated, (1.5 µg/sample, S32354, Invitrogen) in 100 µL SB for 45 minutes at 4 °C followed by three times wash as before. Peptide bound cells were resuspended in 300-500 uL SB and analyzed by BD Accuri C6 Plus Flow Cytometer at medium flow rate for 100 K event. Cells incubated only with Streptavidin-Alexa Fluor 488 were used for background controls. After collecting the data, further analysis was done with FlowJo software where single cells populations were gated and then calculated the percentage of the cells shifted from the background control sample. Cells binding $EC_{50}$ estimated using the GraphPad Prism software.

### Figures

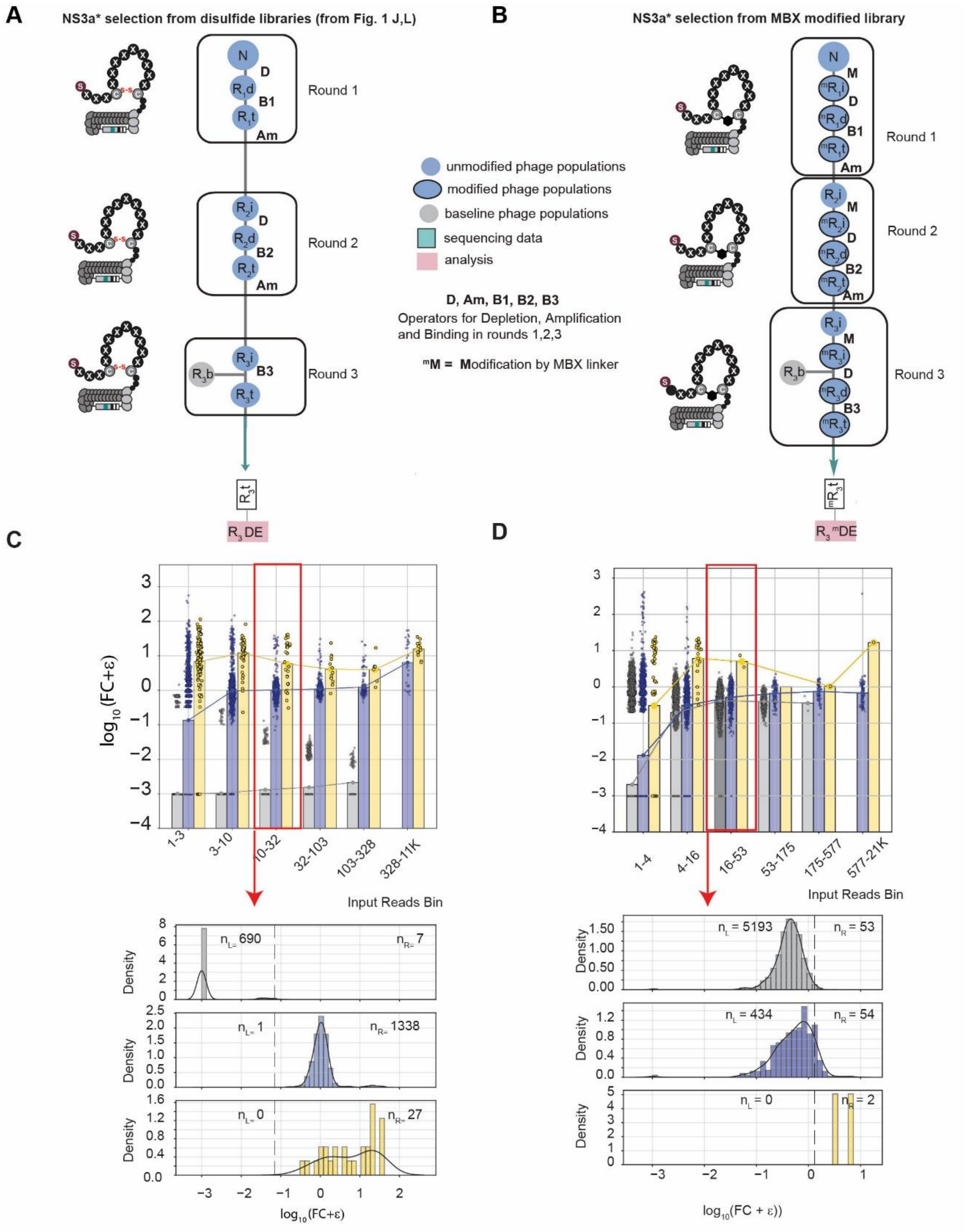

**Fig. S1.** Inserting a baseline library to test the fidelity of .NS3a* selection on disulfide and MBX-modified libraries A) Flow chart of panning campaign against NS3a* with baseline on a disulfide library. B) Flow chart of panning campaign against NS3a* with baseline on

MBX-modified library C) Bar plots showing enrichment of disulfide library after binding by input abundance. The NS3a* selection (blue and yellow bars) shows strong divergence from the non-binding baseline D) Bar plots showing enrichment of MBX-modified library after binding by input abundance.

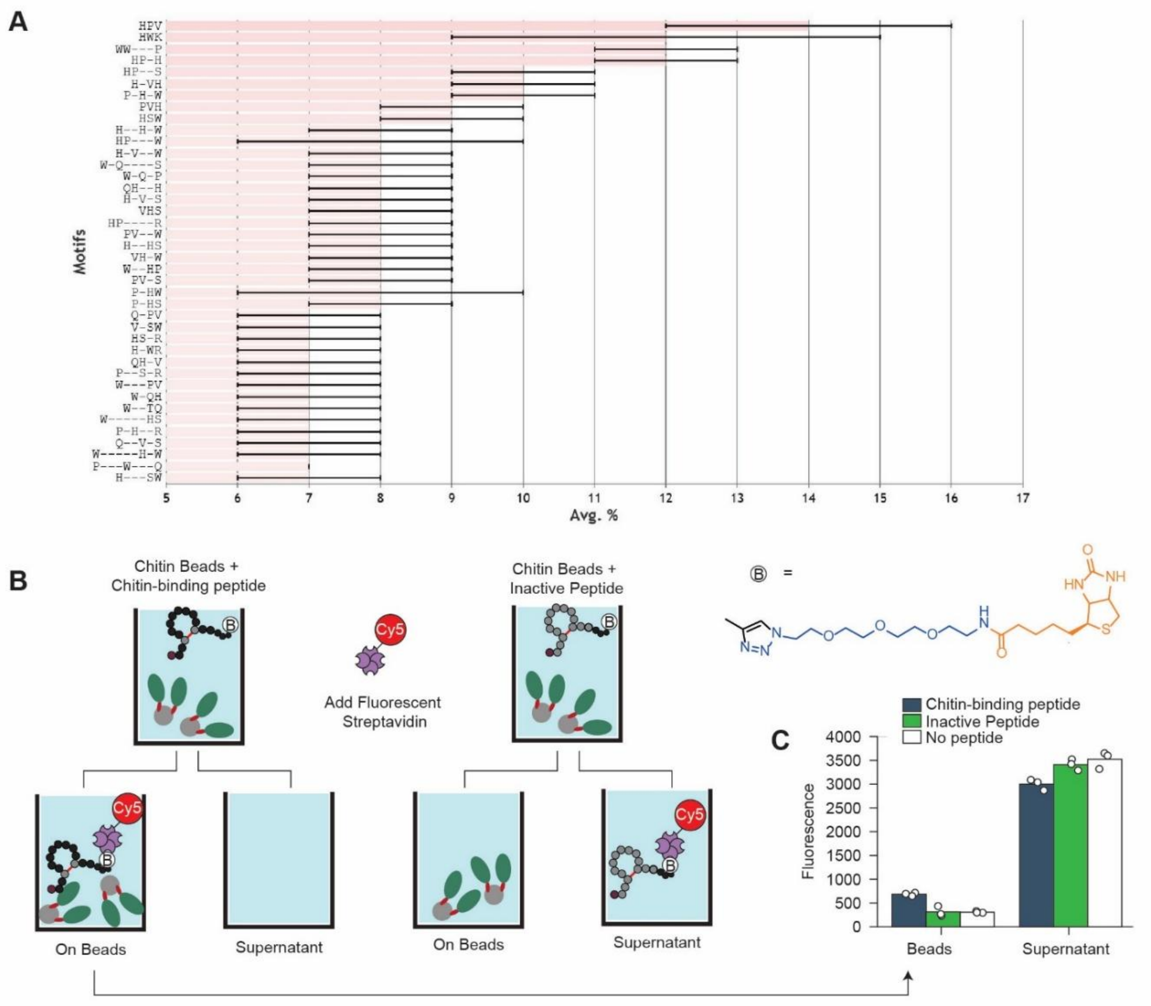

**Fig. S2.** Validation of selection of S2C8C2 library on chitin. A) Motif analysis of round 3 chitin selection output. B) Scheme of procedure for validation of binding of chitin-binding peptide on chitin magnetic beads. C) Binding profile of chitin-binding peptide relative to a random inactive biotinylated peptide sequence.

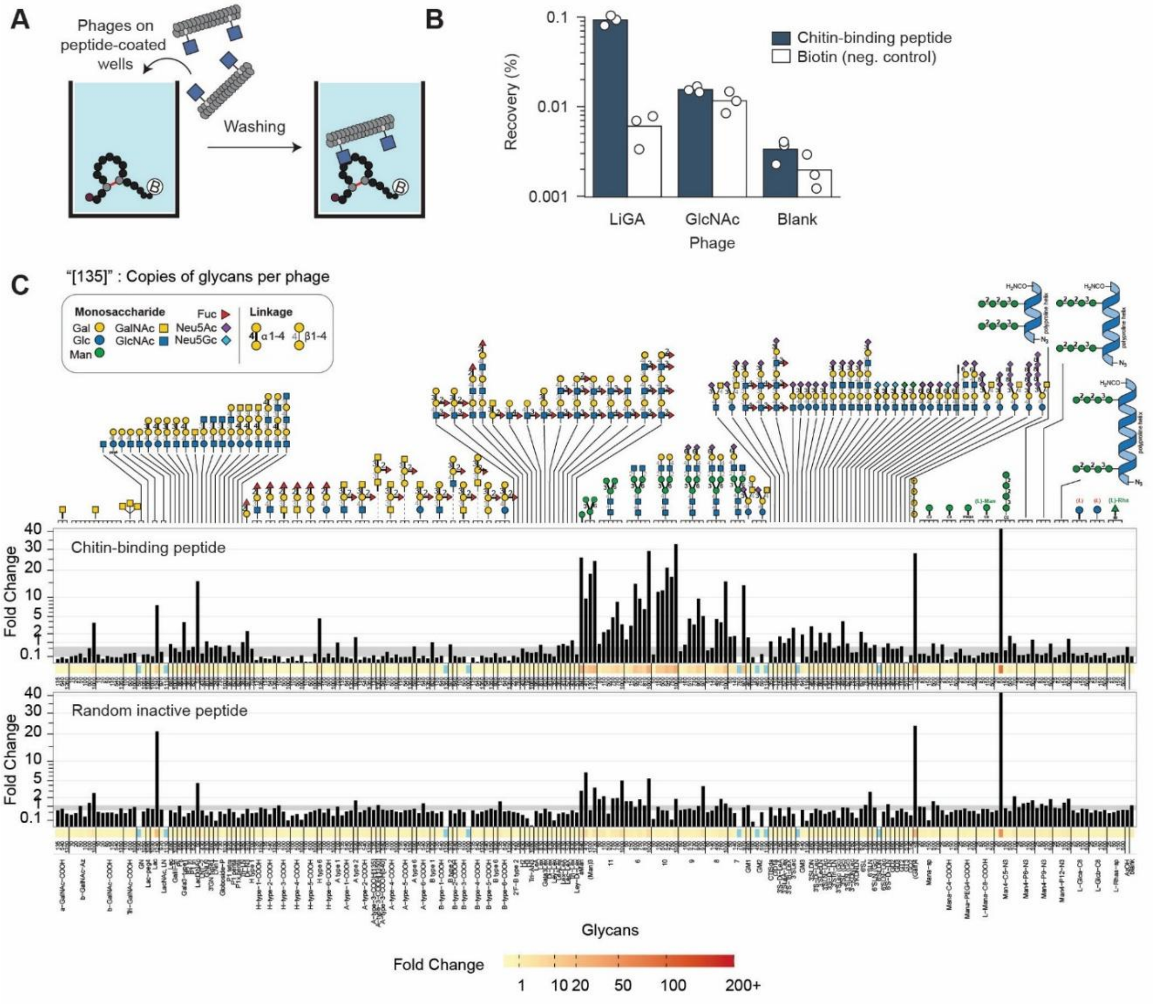

**Figure S3.** Validation of chitin selection using LiGA**.** A) Scheme of procedure for binding of glycan-displaying phages wells coated with biotinylated chitin-binding peptide. B) Recovery of phages from plate panning as measured by qPCR. C) Glycan binding profile of chitin-binding peptide as measured by LiGA.

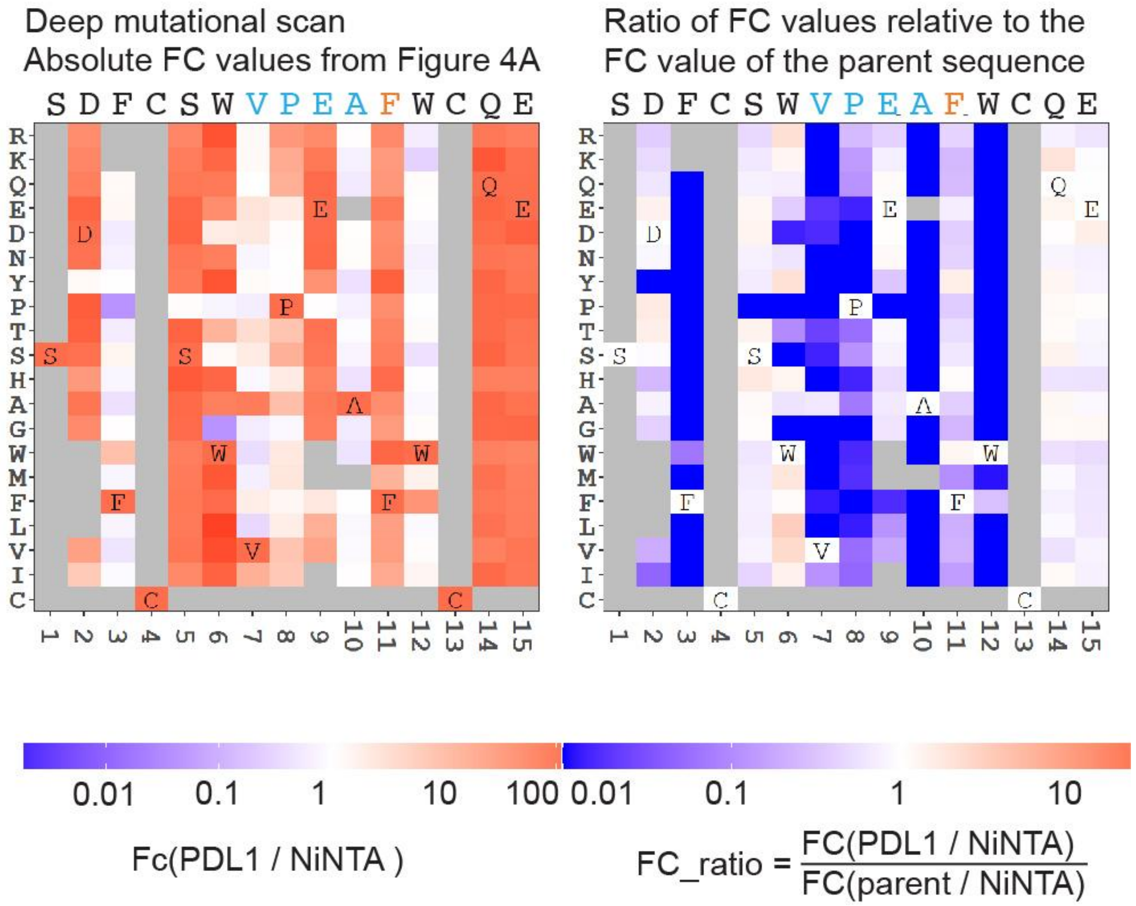

**Figure S4.** Absolute and ratiometric analysis of the FC values depicts in the Main Text Figure 4A. Left panel is the exact copy of the heatmap from Figure 4A describing the enrichment values for all peptides in a deep mutational scan of sequence SDFCSWVPEAFWCQE. On the right, the same heat map has been normalized by calculating the FC_ratio by division by the FC of the SDFCSWVPEAFWCQE sequence highlighting plausible improvements with FC_ratio>1

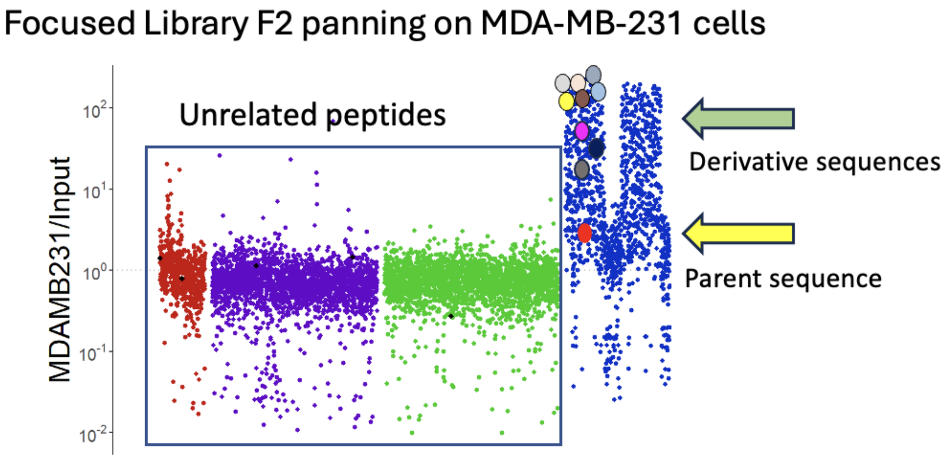

| | Peptide sequences | Input_CPM | MDAMB231_CPM | FC.MDAMB231. vs.Input |
|---|---|---|---|---|
| 8604 | SDFCSLVPEAWWCQEIGG | 15 | 3957 | 271 |
| 8605 | SDFCSLVPEAWWCQETGG | 22 | 5063 | 227 |
| 8606 | STFCSLVPEAWWCQESGG | 15 | 3462 | 226 |
| 8607 | SDFCSLVPEAWWCQEGLG | 47 | 9251 | 195 |
| 8778 | SDFCSLVPEAWWCQEKGG | 23 | 4638 | 205 |
| 8609 | STFCSLVPDAFWCQEGEG | 13 | 1462 | 114 |
| 8404 | SDFCSLVPEAFWCQEGGG | 34 | 322 | 10 |
| 8405 | SDFCSWVPEAYWCQEGGG | 45 | 801 | 18 |
| 8406 | SDFCSWVPEAWWCQEGGG | 14 | 418 | 30 |
| 8191 | SDFCSWVPEAFWCQEGGG | 35 | 109 | 3 |

Parent sequence

**Figure S5.** A broader deep mutational scan for SDFCSWVPEAFWCQEGGG and SDFCSLVPEAFWCQEGGG sequences that include mutations in the S1 and GGG linker regions. The DNA focused library F2 containing these sequences has been cloned into the phage display vector analogously to F1 (Figure 3). The resulting phage displayed library has been panned against PD-L1(+) breast cancer cell line MDA-MB-231. The Manhattan plot describes the FC enrichment of unrelated peptides, which form a baseline response, and the peptides derived from the parent PD-L1-binding sequences. In this phage display campaign the SDFCSWVPEAFWCQEGGG and SDFCSLVPEAFWCQEGGG sequences are only modestly different from the baseline response (i.e., binding of unrelated peptides to MDA-MB-231 cells). On the other hand, many peptides with single amino acid substitution, some of which are illustrated in the table below, exhibit a significantly higher FC value. The binding performance of the synthetic peptide macrocycles corresponding to the parent and the derivative sequences is summarized in Figure 4.

8774:SDFCSLVPEAWWCQEIGG(KB)

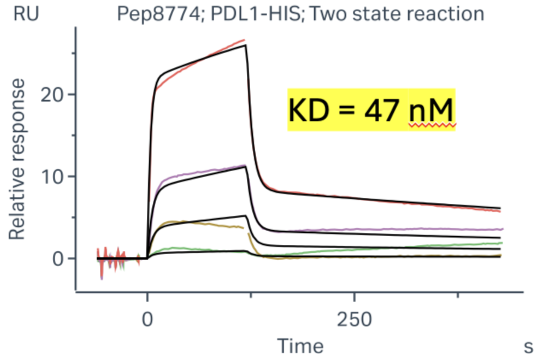

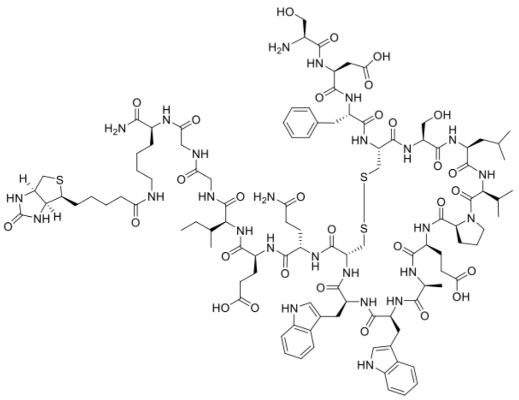

| PDL1-HIS vs Pep 8774 | ka1 (1/Ms) | kd1 (1/s) | ka2 (1/s) | kd2 (1/s) | KD (M) |
|---|---|---|---|---|---|
| Kinetics Model (Two State Reaction) | 6.98e+05 | 1.46e-01 | 3.58e-03 | 1.03e-03 | **4.69e-08** |
| Conc. (nM) | 3.2 | 20 | 50 | 200 | - |

8915:ADFCSLVPEAWWCQEIGGG(KB)

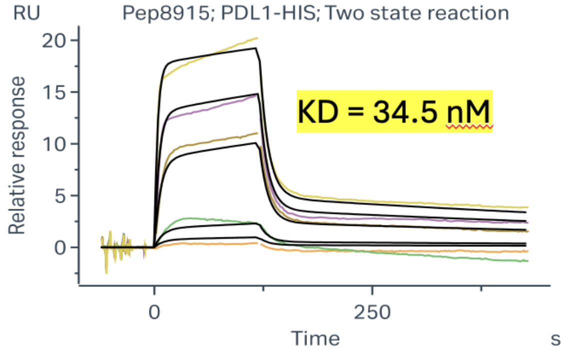

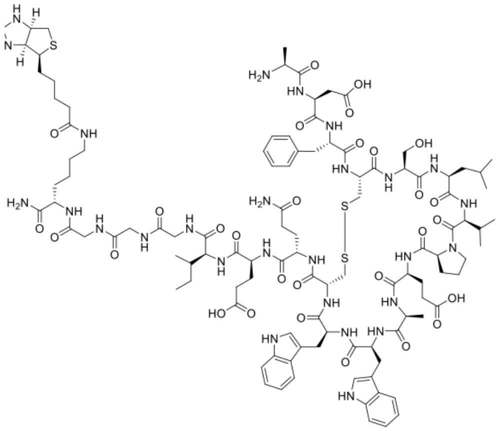

| PDL1-HIS vs Pep 8915 | ka1 (1/Ms) | kd1 (1/s) | ka2 (1/s) | kd2 (1/s) | KD (M) |
|---|---|---|---|---|---|
| Kinetics Model (Two State Reaction) | 9.51e+05 | 1.01e-01 | 2.52e-03 | 1.21e-03 | **3.45e-08** |
| Conc. (nM) | 3.2 | 8 | 50 | 100 | 200 |

**Figure S6.** Representative surface plasmon resonance (SPR) analysis of the binding of macrocyclic peptides 8774 and 8915 (structures displayed) to PD-L1 protein immobilized on the surface of the chip. Kinetic parameters and calculated kinetic Kd are displayed.

9372: ATFCSLVπTAWWCQEG-N(Me)H

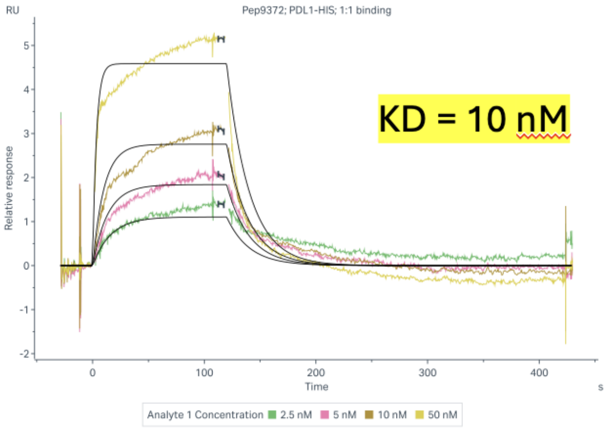

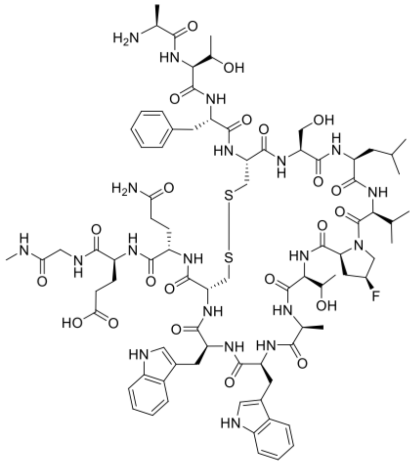

| ka (1/Ms) | kd (1/s) | KD (nM) |
|---|---|---|
| 4.99e+06 | 4.95e-02 | 9.9 |

Steady State KD= 11.9 nM

**Figure S7.** Representative surface plasmon resonance (SPR) analysis of the binding of macrocyclic peptides 9372 (structures displayed) to PD-L1 protein immobilized on the surface of the chip. Kinetic parameters and calculated kinetic Kd are displayed. Steady state Kd was calculated as a dose response of the RUmax.

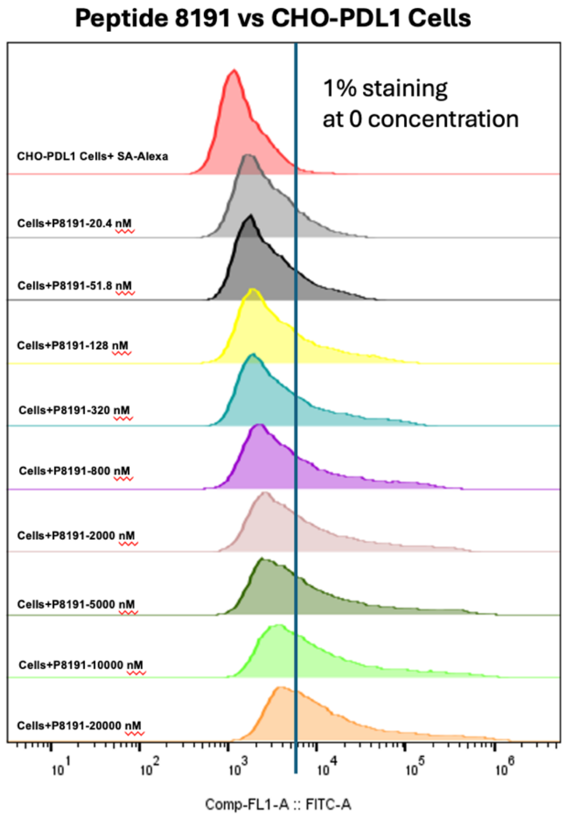

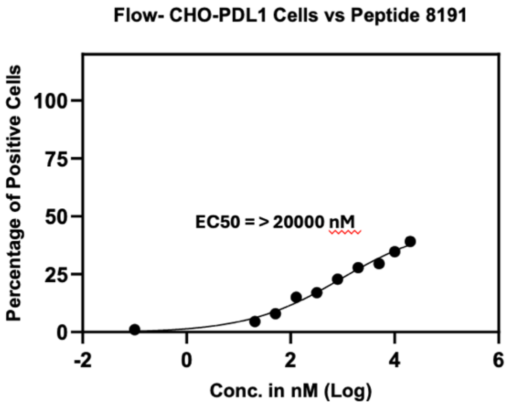

| Peptide Conc. nM | Percentage of Positive Cells with Pep 8191 |
|---|---|
| 0 | 1.07 |
| 20.5 | 4.5 |
| 51.2 | 7.97 |
| 128 | 15 |
| 320 | 17 |
| 800 | 22.9 |
| 2000 | 27.9 |
| 5000 | 29.5 |
| 10000 | 34.8 |
| 20000 | 39.1 |

**Figure S8.** Representative flow-cytometry analysis of the dose-response titration of biotinylated macrocyclic peptides 8191 to PD-L1(+) HEK cells, as measured by secondary reporter, Alexa488-streptavidin. The EC50 was calculated by estimating the concentration at which 50% of the cell population is stained by the biotinylated macrocycle + Alexa488-streptavidin (here, not attained even at the highest concentration of peptide).

**Pep8774:**SS-SDFCSLVPEAWWCQEIGG{KB}

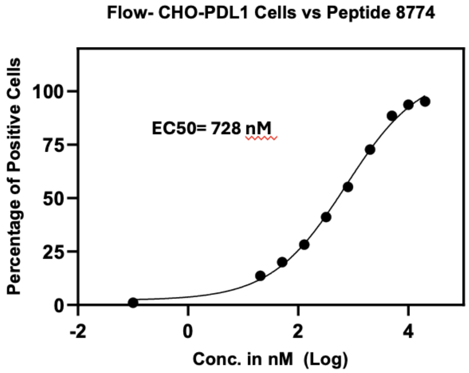

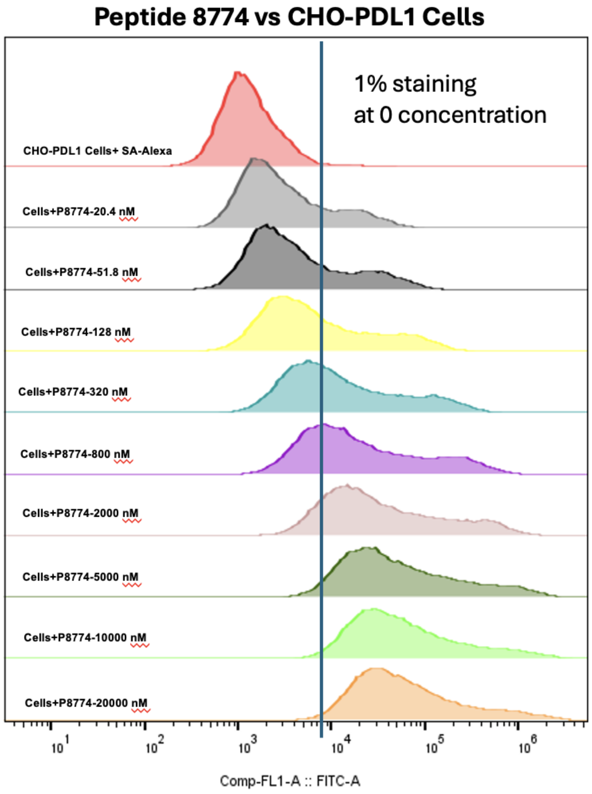

| log(agonist) vs. response -- Variable slope (four parameters) | Pep8774 |
|---|---|
| Best-fit values | |
| Bottom | 2.178 |
| Top | 109.4 |
| LogEC50 | 2.862 |
| HillSlope | 0.6418 |
| EC50 | 728.2 |
| Span | 107.2 |
| 95% CI (profile likelihood) | |
| Bottom | -3.907 to 7.737 |
| Top | 99.68 to 125.7 |
| LogEC50 | 2.692 to 3.105 |
| HillSlope | 0.4960 to 0.8184 |
| EC50 | 492.4 to 1273 |
| Goodness of Fit | |
| Degrees of Freedom | 6 |
| R squared | 0.9972 |
| Sum of Squares | 31.43 |
| Sy.x | 2.289 |
| Number of points | |
| # of X values | 10 |
| # Y values analyzed | 10 |

**Figure S9.** Representative flow-cytometry analysis of the dose-response titration of biotinylated macrocyclic peptides 8774 (structure in **Fig. S6**) to PD-L1(+) HEK cells, as measured by secondary reporter, Alexa488-streptavidine. The EC50 was calculated by estimating the concentration at which 50% of the cell population is stained by the biotinylated macrocycle + Alexa488-streptavidin (here, attained at ~700 nM concentration).

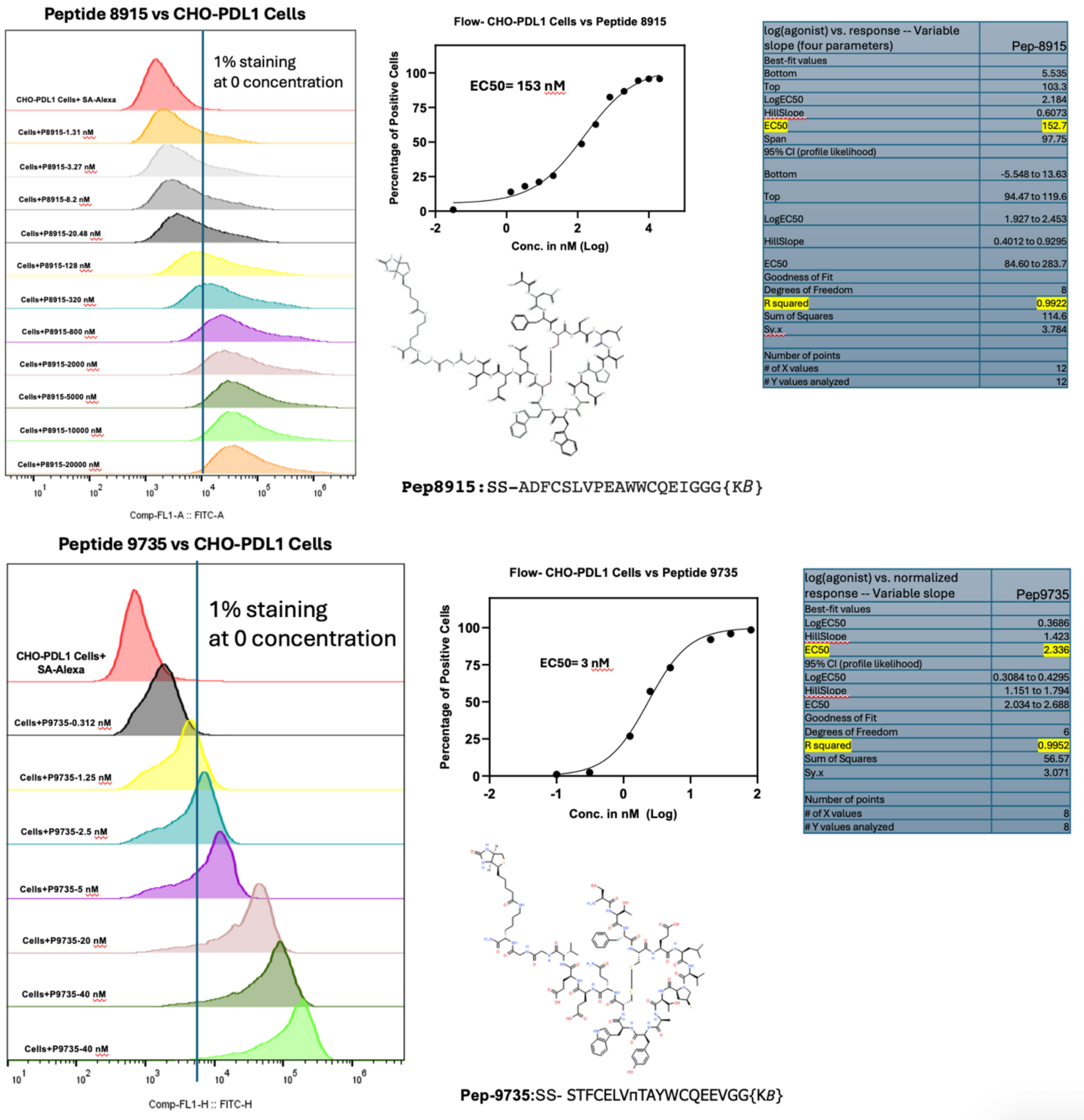

**Figure S10.** Representative flow-cytometry analysis of the dose-response titration of biotinylated macrocyclic peptides 8915 and 9775 (structure shown) to PD-L1(+) HEK cells, as measured by secondary reporter, Alexa488-streptavidin. The EC50 was calculated by estimating the concentration at which 50% of the cell population is stained by the biotinylated macrocycle + Alexa488-streptavidine. Note that in case of peptide 9735, the estimate of EC50=3 nM is conservative and the EC50 value ranges from 300 pM to 5 nM to depends on the threshold established for the non-stained population.

Peptide 8191:
DS-SDFCSWVPEAFWCQEGGG{*KB*}

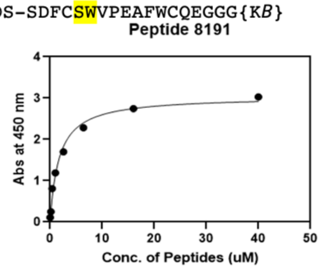

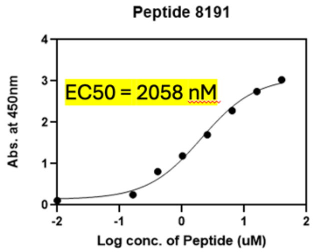

| One site -- Specific binding | |
|---|---|
| **Best-fit values** | |
| Bmax | 3.040 |
| **Kd** | **1.708 uM** |
| **95% CI (profile likelihood)** | |
| Bmax | 2.770 to 3.337 |
| Kd | 1.156 to 2.505 |
| **Goodness of Fit** | |
| Degrees of Freedom | 6 |
| R squared | 0.9878 |
| Sum of Squares | 0.1050 |
| Sy.x | 0.1323 |
| **Number of points** | |
| # of X values | 8 |
| # Y values analyzed | 8 |

| Conc. of Peptides (uM) | Pep8191 |
|---|---|
| 40 | 3.0279 |
| 16 | 2.7458 |
| 6.4 | 2.2826 |
| 2.56 | 1.6996 |
| 1.024 | 1.1900 |
| 0.4096 | 0.8091 |
| 0.16384 | 0.2477 |
| 0 | 0.1077 |

Pep8774:
SS-SDFCSLVPEAWWCQEIGG{KB}

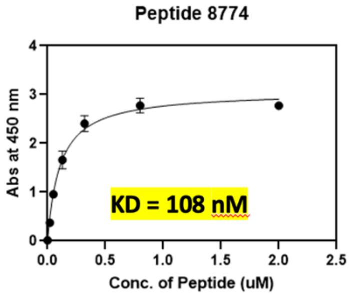

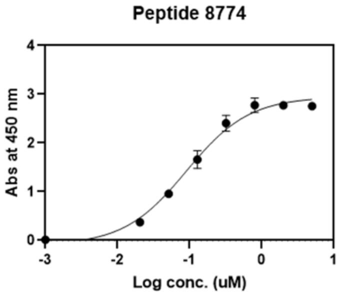

| Log conc. (uM) | Pep8774 | Pep8774 |
|---|---|---|
| 0.698970004 | 2.7218 | 2.7883 |
| 0.301029996 | 2.8136 | 2.7314 |
| -0.096910013 | 2.6707 | 2.8786 |
| -0.494850022 | 2.2884 | 2.5217 |
| -0.89279003 | 1.5271 | 1.7881 |
| -1.290730039 | 0.9503 | 0.9566 |
| -1.688670048 | 0.3414 | 0.4002 |
| 0 | 0.0127 | 0.0114 |

**Figure S11.** Representative dose-response titrations read out by ELISA for two macrocyclic peptides with C-terminal biotin (8191 and 8774, sequences shown, KB denotes the C-terminal biotin-lysine, see Figure S10 for the representative structure). Peptides were incubated with PD-L1 coated plates, the plates were washed and probed with Streptavidin-HRP

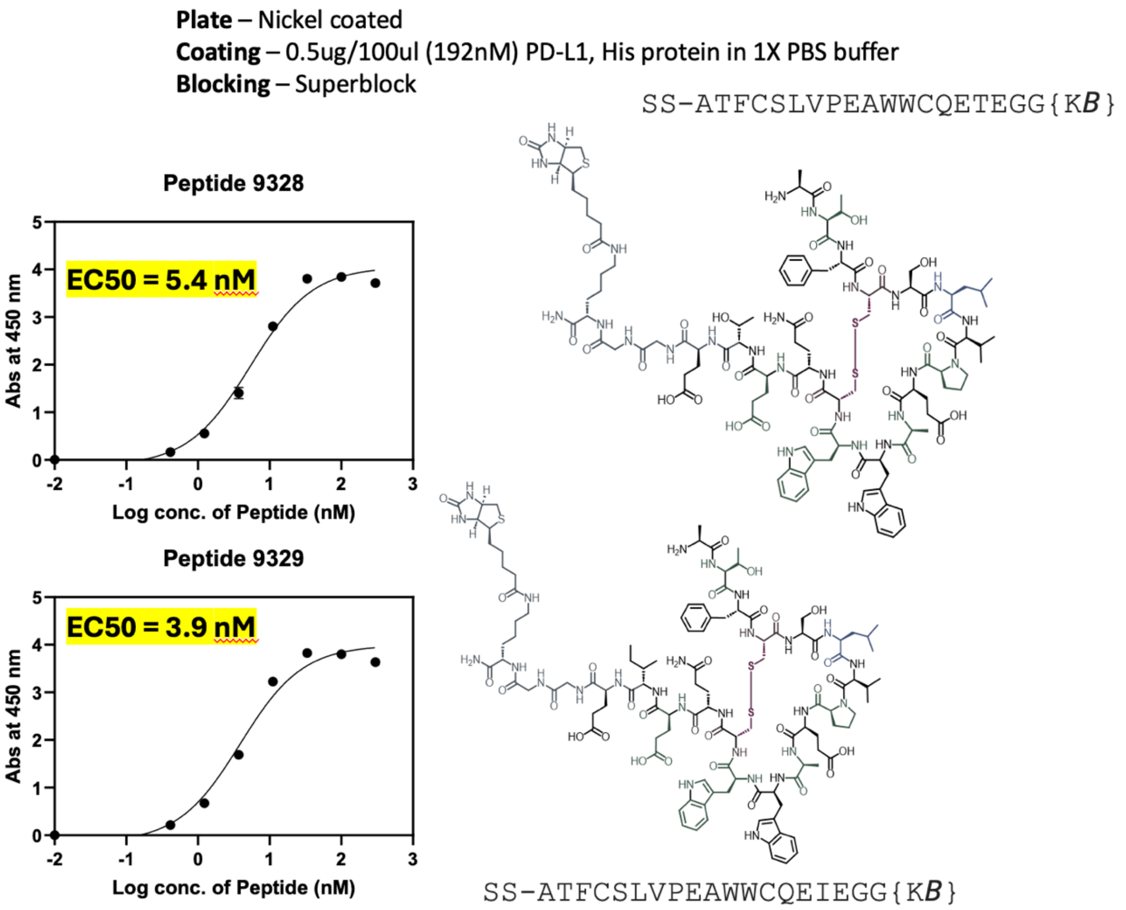

**Figure S12.** Representative dose-response titrations read out by ELISA for two optimized macrocyclic peptides with C-terminal biotin (sequences and structures shown). Peptides were incubated with PD-L1 coated plates, the plates were washed and probed with Streptavidin-HRP

### Tables

**Table S1.** Pre-selection NGS files with unmodified disulfide libraries.

| Library | R1 Input | R1 Output | R2 Input | R2 Output |
|---|---|---|---|---|
| $SX_2CX_8CX_2$ | https://48hd.cloud/file/14384 | https://48hd.cloud/file/14381 | https://48hd.cloud/file/14386 | https://48hd.cloud/file/14383 |
| $SX_3CX_9C$ | https://48hd.cloud/file/14391 | https://48hd.cloud/file/14388 | https://48hd.cloud/file/14393 | https://48hd.cloud/file/14390 |

**Table S2.** NS3a*-selection panning NGS files.

| | Library | R3 Input | R3 Output | Control |
|---|---|---|---|---|
| With baseline | SX3CX9C | https://48hd.cloud/file/16919 | https://48hd.cloud/file/16918 | https://48hd.cloud/file/16920 |
| Without baseline | SX3CX9C | https://48hd.cloud/file/16426 | https://48hd.cloud/file/16425 | https://48hd.cloud/file/16516 |
| With baseline | $SX_3CX_9C$ | https://48hd.cloud/file/17663 | https://48hd.cloud/file/17662 | - |

**Table S3.** Chitin-selection panning NGS files.

| Library | R3 Input | R3 Output |
|---|---|---|
| $SX_2CX_8CX_2$ + GS23 | http://48hd.cloud/file/19871 | https://48hd.cloud/file/19870 |

**Table S4.** Chitin-selection panning mixed with NS3a* library NGS files.

| Library | R3 Input | R3 Output |
|---|---|---|
| $SX_2CX_8CX_2$ + $SX_3CX_9C$ + GS23 | https://48hd.cloud/file/21525 | https://48hd.cloud/file/21527 |

**Table S5.** NS3a*-selection panning mixed with chitin library NGS files.

| Library | R3 Input | R3 Output |
|---|---|---|
| $SX_2CX_8CX_2$ + $SX_3CX_9C$ + GS23 | https://48hd.cloud/file/21525 | https://48hd.cloud/file/21522 |
