## Supplementary material for "Universal Baseline for *in vitro* Selection of Genetically Encoded Libraries": Dataset: bin_bar_chart.py

```
"""
Complete visualization and analysis functions for peptide NGS data.

This module provides functions for loading, analyzing, and visualizing peptide NGS reads data,
including separate bin and scatter plots with motif highlighting and group selection.

Author: Kejia
Date: May 08, 2025
Modified: May 20, 2025 - Added support for custom motifs and group selection
"""
import pandas as pd
import numpy as np
import matplotlib.pyplot as plt
import seaborn as sns
import matplotlib.patheffects as path_effects
import matplotlib.ticker as ticker
import re


# ============================================================================
# DATA LOADING AND ANALYSIS FUNCTIONS
# ============================================================================

def load_data(file_path, min_cpm=None):
    """
    Load and validate the peptide NGS reads data.
    
    Parameters:
    -----------
    file_path : str
        Path to the CSV file containing peptide data
    min_cpm : float, optional
        Minimum counts per million to include
        
    Returns:
    --------
    pandas.DataFrame or None
        The loaded dataframe, or None if loading failed
    """
    try:
        df = pd.read_csv(file_path)
        print(f"Successfully loaded data with {len(df)} rows and {len(df.columns)} columns.")
        
        if min_cpm is not None:
            original_count = len(df)
            df = df[df["input_average"] >= min_cpm]
            filtered_count = len(df)
            print(f"Filtered data using minimum CPM of {min_cpm}: removed {original_count - filtered_count} rows, {filtered_count} rows remaining")
        
        return df
    except Exception as e:
        print(f"Error loading file: {e}")
        return None


def identify_motif_sequences(df, motif, sequence_col="sequence"):
    """
    Identify sequences containing a specific motif.
    
    Parameters:
    -----------
    df : pandas.DataFrame
        The dataframe containing peptide sequences
    motif : str
        The motif to search for
    sequence_col : str, default="sequence"
        Column name for peptide sequences
        
    Returns:
    --------
    numpy.ndarray
        Boolean mask indicating which sequences contain the motif
    """
    if sequence_col not in df.columns:
        print(f"Warning: {sequence_col} column not found in dataframe")
        return np.zeros(len(df), dtype=bool)
    
    has_motif = df[sequence_col].str.contains(motif, case=False, na=False)
    print(f"Found {has_motif.sum()} sequences containing '{motif}'")
    return has_motif


def create_simplified_bin_ranges(min_val, max_val, num_bins=8):
    """
    Create simplified bin ranges and labels with pretty formatting.
    
    Parameters:
    -----------
    min_val : float
        Minimum value for binning
    max_val : float
        Maximum value for binning
    num_bins : int, default=8
        Number of bins to create
        
    Returns:
    --------
    tuple
        (bins, bin_labels) - List of bin edges and list of formatted bin labels
    """
    if min_val <= 0:
        min_val = 1
    
    # Use logarithmic binning for better spread
    log_bins = np.logspace(np.log10(min_val), np.log10(max_val), num_bins+1)
    bins = log_bins.tolist()
    
    # Create simplified labels
    bin_labels = []
    for i in range(len(bins)-1):
        start = int(bins[i])
        end = int(bins[i+1])
        
        # Format numbers with K and M suffixes for readability
        if start >= 1000000:
            start_str = f"{start/1000000:.0f}M"
        elif start >= 1000:
            start_str = f"{start/1000:.0f}K"
        else:
            start_str = f"{start}"
            
        if end >= 1000000:
            end_str = f"{end/1000000:.0f}M"
        elif end >= 1000:
            end_str = f"{end/1000:.0f}K"
        else:
            end_str = f"{end}"
            
        bin_labels.append(f"{start_str}-{end_str}")
    
    return bins, bin_labels


def process_data_bins(df, input_col="input_average", fc_col="FC", group_col="group", tail_merge_threshold=None):
    """
    Process data into bins for bar plots.
    
    Parameters:
    -----------
    df : pandas.DataFrame
        The original data frame with all peptide data
    input_col : str, default="input_average"
        Column name for input reads
    fc_col : str, default="FC"
        Column name for fold change values
    group_col : str, default="group"
        Column name for group designation
        
    Returns:
    --------
    tuple
        (input_bins, input_bin_labels, input_binned_df)
    """
    # Find min and max input values
    min_input = max(df[input_col].min(), 1)
    max_input = df[input_col].max()
    
    # Define simplified bin ranges for input plot
    input_bins, input_bin_labels = create_simplified_bin_ranges(min_input, max_input, num_bins=8)
    print("Initial bins:", input_bins)
    print("Initial labels:", input_bin_labels)
    
    # Optional: merge all bins whose *lower edge* is strictly above tail_merge_threshold
    if tail_merge_threshold is not None:
        tail_start_idx = None

        # find first bin whose lower edge is strictly > threshold
        for i in range(len(input_bins) - 1):
            if input_bins[i] > tail_merge_threshold:
                tail_start_idx = i
                break

        if tail_start_idx is not None:
            # How many tail bins would be merged?
            # (bins are intervals [input_bins[k], input_bins[k+1]))
            num_tail_bins = (len(input_bins) - 1) - tail_start_idx

            # Only merge if there are at least *two* bins in the tail
            if num_tail_bins >= 2:
                merged_bins = input_bins[:tail_start_idx + 1] + [input_bins[-1]]

                # Rebuild labels so they match the merged bins
                merged_labels = []
                for j in range(len(merged_bins) - 1):
                    start = int(merged_bins[j])
                    end = int(merged_bins[j + 1])

                    if start >= 1_000_000:
                        start_str = f"{start/1_000_000:.0f}M"
                    elif start >= 1_000:
                        start_str = f"{start/1_000:.0f}K"
                    else:
                        start_str = f"{start}"

                    if end >= 1_000_000:
                        end_str = f"{end/1_000_000:.0f}M"
                    elif end >= 1_000:
                        end_str = f"{end/1_000:.0f}K"
                    else:
                        end_str = f"{end}"

                    merged_labels.append(f"{start_str}-{end_str}")

                print("Initial bins:", input_bins)
                print("Initial labels:", input_bin_labels)
                input_bins = merged_bins
                input_bin_labels = merged_labels
                print(f"Merged tail bins above {tail_merge_threshold} into a single bin: {merged_labels[-1]}")
                print("Final bins after merge:", input_bins)
                print("Final labels after merge:", input_bin_labels)
    # Get unique groups
    if group_col in df.columns:
        groups = sorted(df[group_col].unique())
    else:
        groups = [1]  # Default group
    
    # Process data by bins for input plot
    input_binned_data = []
    
    for i in range(len(input_bins)-1):
        bin_start = input_bins[i]
        bin_end = input_bins[i+1]
        bin_center = (bin_start + bin_end) / 2
        
        for group in groups:
            if group_col in df.columns:
                group_data = df[df[group_col] == group]
            else:
                group_data = df
            
            # Get data points in this bin
            bin_data = group_data[(group_data[input_col] >= bin_start) & 
                                (group_data[input_col] < bin_end)]
            
            if len(bin_data) > 0:
                avg_fc = bin_data[fc_col].mean()
                
                input_binned_data.append({
                    'bin_label': input_bin_labels[i],
                    'bin_index': i,
                    'bin_center': bin_center,
                    'bin_start': bin_start,
                    'bin_end': bin_end,
                    'group': group,
                    'avg_fc': avg_fc,
                    'count': len(bin_data)
                })
    
    # Convert to DataFrame for easier plotting
    input_binned_df = pd.DataFrame(input_binned_data)
    
    return input_bins, input_bin_labels, input_binned_df


def check_available_groups(df, group_col="group"):
    """Helper function to see what groups are available in your dataset"""
    if group_col in df.columns:
        groups = sorted(df[group_col].unique())
        print(f"Available groups in your dataset: {groups}")
        for group in groups:
            count = (df[group_col] == group).sum()
            print(f"  Group {group}: {count} sequences")
        return groups
    else:
        print(f"Column '{group_col}' not found in dataframe")
        return []


# ============================================================================
# STEP 2: Replace your plot_binned_bar_chart function with this one
# ============================================================================

def plot_binned_bar_chart(df, input_binned_df, input_bins, input_bin_labels, has_motif=None,
                         output=None, y_scale="log", max_fc=None, 
                         input_col="input_average", fc_col="FC", group_col="group", sequence_col="sequence", 
                         y_min=None, x_max=None, motif="DMT", custom_motif=None, 
                         groups_to_plot=None, group_names=None, colors=None, used_bins=None,
                         show_motifs=True, transform_data="none", min_bar_height_ratio=0.33, 
                         baseline_group=None, motif_group=None):
    """
    Create a binned bar chart of FC values by input reads.
    Lines will automatically connect the averages of each bin.
    
    Parameters:
    -----------
    transform_data : str, default="none"
        Data transformation: "none", "sqrt", "log"
        - "none": No transformation
        - "sqrt": Apply sqrt(data) then plot with y_scale
        - "log": Apply log(data) then plot with y_scale
    min_bar_height_ratio : float, default=0.33
        Position of baseline group mean as a fraction of the y-axis range (0.33 = 1/3 of plot).
        The y-axis will be scaled so that the mean of ONLY baseline group bars appears at this height.
        Library and motif bars will scale proportionally, preserving their relative differences.
        For example, 0.33 means the baseline mean appears at 1/3 from the bottom.
        Set to 0 or None to disable automatic y-axis scaling.
    baseline_group : int, optional
        Which group number to use as baseline for y-axis scaling. If None, uses the lowest
        numbered group. Only bars from this group are used to calculate the mean for positioning.
    """
    # Filter dataframe to only include specified groups
    if groups_to_plot is not None:
        df_filtered = df[df[group_col].isin(groups_to_plot)].copy()
        input_binned_df_filtered = input_binned_df[input_binned_df['group'].isin(groups_to_plot)].copy()
        print(f"Filtering to show only groups: {groups_to_plot}")
    else:
        df_filtered = df.copy()
        input_binned_df_filtered = input_binned_df.copy()
        groups_to_plot = sorted(df[group_col].unique()) if group_col in df.columns else [1]
    
    # Create figure and axis
    fig, ax = plt.subplots(figsize=(8, 8))
    
    # Define color palette for groups
    if group_col in df_filtered.columns:
        groups = sorted(df_filtered[group_col].unique())
        
        # Use provided colors or create default color mapping
        if colors is None:
            all_possible_groups = groups_to_plot if groups_to_plot else groups
            palette = sns.color_palette("husl", len(all_possible_groups))
            colors = {g: palette[i] for i, g in enumerate(all_possible_groups)}
        
        # Use provided group names or create default names
        if group_names is None:
            group_names = {g: f"Group {g}" for g in groups}
    else:
        groups = [1]  # Default group
        colors = {1: 'blue'} if colors is None else colors
        group_names = {1: 'All Data'} if group_names is None else group_names
    
    # Count zero values
    zero_count = (df_filtered[fc_col] == 0).sum()
    if zero_count > 0:
        print(f"Found {zero_count} points with FC = 0")
    
    # Apply data transformation if requested
    df_plot = df_filtered.copy()
    y_label_suffix = ""
    
    if transform_data == "sqrt":
        print("Applying square root transformation to data")
        # Handle negative values and zeros
        min_val = df_plot[fc_col].min()
        if min_val <= 0:
            shift = abs(min_val) + 0.001
            print(f"Shifting data by +{shift:.3f} to make all values positive for sqrt transformation")
            df_plot[fc_col] = df_plot[fc_col] + shift
        
        df_plot[fc_col] = np.sqrt(df_plot[fc_col])
        y_label_suffix = " (sqrt transformed)"
        
    elif transform_data == "log":
        print("Applying log transformation to data")
        # Handle negative values and zeros
        min_val = df_plot[fc_col].min()
        if min_val <= 0:
            shift = abs(min_val) + 0.001
            print(f"Shifting data by +{shift:.3f} to make all values positive for log transformation")
            df_plot[fc_col] = df_plot[fc_col] + shift
        
        df_plot[fc_col] = np.log10(df_plot[fc_col])
        y_label_suffix = " (log10 transformed)"
    elif transform_data == "log_eps":
        # Log10(FC + eps) for linear y-axis
        eps = 1e-3
        print(f"Applying log10({fc_col} + {eps}) transformation to data (for linear y-axis)")
        
        fc_vals = df_plot[fc_col].astype(float).values
        
        # Clamp negatives to zero to avoid weird logs
        fc_vals = np.where(fc_vals < 0, 0, fc_vals)
        
        df_plot[fc_col] = np.log10(fc_vals + eps)
        y_label_suffix = f" (log10({fc_col} + {eps}))"
    
    # Recalculate zero count for transformed data
    zero_count = (df_plot[fc_col] == 0).sum()
    if zero_count > 0:
        print(f"Found {zero_count} points with FC = 0 after transformation")
    
    # Only process motifs if show_motifs is True
    has_custom_motif = None
    if show_motifs and custom_motif is not None and sequence_col in df_plot.columns:
        pattern = re.compile(custom_motif)
        has_custom_motif = df_plot[sequence_col].apply(lambda x: bool(pattern.search(x)) if isinstance(x, str) else False)
        motif_count = has_custom_motif.sum()
        print(f"Binned Bar Chart: Found {motif_count} sequences containing the custom motif '{custom_motif}'")
    
    # Print primary motif info only if show_motifs is True
    if show_motifs and has_motif is not None:
        # Filter has_motif to match the filtered dataframe
        has_motif_filtered = has_motif.loc[df_plot.index] if hasattr(has_motif, 'loc') else has_motif
        print(f"\nBinned Bar Chart: Found {has_motif_filtered.sum()} {motif} sequences")
        if group_col in df_plot.columns:
            for g in sorted(df_plot[group_col].unique()):
                g_motif_count = ((df_plot[group_col] == g) & has_motif_filtered).sum()
                print(f"Group {g}: {g_motif_count} {motif} sequences")
    else:
        has_motif_filtered = None
    
    # Calculate the FC range for y-axis using transformed data
    positive_fc = df_plot[df_plot[fc_col] > 0][fc_col]
    
    if len(positive_fc) > 0:
        fc_min = min(0.01, positive_fc.min() * 0.8)
    else:
        fc_min = 0.01
        print("Warning: No positive FC values found for log scale calculations")
    
    if max_fc:
        fc_max = max_fc
    else:
        fc_max = max(10.0, df_plot[fc_col].max() * 1.1)
    
    # Calculate y-axis limits to position BASELINE mean at 1/3 of plot height
    baseline_position_ratio = None
    adjusted_y_max = None
    
    if min_bar_height_ratio and min_bar_height_ratio > 0:
        # Calculate mean of ONLY baseline bars (exclude library and motif bars)
        baseline_bar_values = []
        
        # Determine which group is baseline
        if baseline_group is not None:
            # Use user-specified baseline group
            baseline_group_id = baseline_group
            print(f"Using user-specified group {baseline_group_id} as baseline for y-axis scaling")
        elif len(groups) > 1:
            # Auto-detect: assume baseline is the lowest numbered group
            baseline_group_id = min(groups)
            print(f"Auto-detected group {baseline_group_id} as baseline for y-axis scaling")
        else:
            # Only one group
            baseline_group_id = groups[0]
            print(f"Using group {baseline_group_id} as baseline for y-axis scaling")
        
        # Get only baseline group data
        baseline_group_data = input_binned_df_filtered[input_binned_df_filtered['group'] == baseline_group_id]
        if len(baseline_group_data) > 0:
            baseline_bar_values = baseline_group_data['avg_fc'].values
        
        if len(baseline_bar_values) > 0:
            mean_baseline = np.mean(baseline_bar_values)
            baseline_position_ratio = min_bar_height_ratio
            
            if y_scale == "log":
                # For log scale: mean_baseline should appear at baseline_position_ratio of the log range
                plot_y_min = y_min if y_min is not None else fc_min
                
                if mean_baseline > plot_y_min:
                    log_mean = np.log10(mean_baseline)
                    log_min = np.log10(plot_y_min)
                    log_max_needed = (log_mean - log_min) / baseline_position_ratio + log_min
                    adjusted_y_max = 10 ** log_max_needed
                    
                    print(f"Baseline group mean = {mean_baseline:.3f}")
                    print(f"Adjusting y-max to {adjusted_y_max:.3f} so baseline mean appears at {baseline_position_ratio*100:.0f}% of plot height")
                elif mean_baseline <= plot_y_min:
                    plot_y_min = mean_baseline / 10.0
                    log_mean = np.log10(mean_baseline)
                    log_min = np.log10(plot_y_min)
                    log_max_needed = (log_mean - log_min) / baseline_position_ratio + log_min
                    adjusted_y_max = 10 ** log_max_needed
            else:
                # For linear scale: mean_baseline = y_min + baseline_position_ratio × (y_max - y_min)
                plot_y_min = 0  # Linear scale typically starts at 0
                
                if mean_baseline > plot_y_min:
                    adjusted_y_max = (mean_baseline - plot_y_min) / baseline_position_ratio + plot_y_min
                    
                    print(f"Baseline group mean = {mean_baseline:.3f}")
                    print(f"Adjusting y-max to {adjusted_y_max:.3f} so baseline mean appears at {baseline_position_ratio*100:.0f}% of plot height")
    
    # Plot binned bar chart
    if len(input_binned_df_filtered) > 0:
        # Calculate positions for grouped bars
        bar_width = 0.25
        if used_bins is None:
            unique_bins = sorted(input_binned_df_filtered['bin_index'].unique())
        else:
            unique_bins = used_bins
        
        # Set up positions for the bars
        x_positions = np.arange(len(unique_bins))
        
        # Create a custom group order
        plot_groups = groups_to_plot
        
        # Plot the regular groups (without motif peptides if motifs are being shown)
        for i, group in enumerate(plot_groups):
            group_data = input_binned_df_filtered[input_binned_df_filtered['group'] == group]
            
            avg_fc_values = []
            bin_data_points = []
            
            for bin_idx in unique_bins:
                bin_group_data = group_data[group_data['bin_index'] == bin_idx]
                if len(bin_group_data) > 0:
                    # For groups when showing motifs, exclude motifs from the average
                    if show_motifs and sequence_col in df_plot.columns:
                        bin_start = input_bins[bin_idx]
                        bin_end = input_bins[bin_idx+1]
                        
                        points_data = df_plot[(df_plot[group_col] == group) & 
                                      (df_plot[input_col] >= bin_start) & 
                                      (df_plot[input_col] < bin_end)]
                        
                        # Exclude motif points
                        exclude_mask = np.zeros(len(points_data), dtype=bool)
                        if has_motif_filtered is not None:
                            exclude_mask |= has_motif_filtered.loc[points_data.index]
                        if has_custom_motif is not None:
                            exclude_mask |= has_custom_motif.loc[points_data.index]
                        
                        non_motif_points = points_data[~exclude_mask]
                        
                        if len(non_motif_points) > 0:
                            avg_fc = non_motif_points[fc_col].mean()
                        else:
                            avg_fc = 0
                        
                        avg_fc_values.append(avg_fc)
                        bin_data_points.append({
                            'bin_index': bin_idx,
                            'x_pos': bin_idx,
                            'points': non_motif_points
                        })
                    else:
                        bin_start = input_bins[bin_idx]
                        bin_end = input_bins[bin_idx+1]
                        points_data = df_plot[
                            (df_plot[group_col] == group) &
                            (df_plot[input_col] >= bin_start) &
                            (df_plot[input_col] < bin_end)
                            ]
                        if transform_data == "sqrt":
                            avg_fc = points_data[fc_col].mean()
                        elif transform_data in ("log", "log_eps"):
                            avg_fc = points_data[fc_col].mean()
                        else:
                            avg_fc = bin_group_data['avg_fc'].values[0]
                        avg_fc_values.append(avg_fc)
                        bin_data_points.append({
                            'bin_index': bin_idx,
                            'x_pos': bin_idx,
                            'points': points_data
                            })
                else:
                    avg_fc_values.append(np.nan)
                    bin_data_points.append(None)
            
            # Position offset for bars
            offset = (i - len(plot_groups)/2) * bar_width
                        # For log_eps: force bars to start from -4
            if transform_data == "log_eps":
                forced_ymin = -4
                bar_bottom = forced_ymin
                bar_heights = [v - forced_ymin for v in avg_fc_values]

                bars = ax.bar(
                    x_positions + offset,
                    bar_heights,
                    width=bar_width,
                    bottom=bar_bottom,
                    #facecolor='none',
                    color=colors[group],
                    #edgecolor=colors[group],   # silhouette color
                    edgecolor='black',
                    linewidth=0.5  
                )
            else:
                bars = ax.bar(
                    x_positions + offset,
                    avg_fc_values,
                    width=bar_width,
                    alpha=0.8,
                    color=colors[group]
                )

            
            # Overlay individual FC points on top of each bar
            for j, bin_data in enumerate(bin_data_points):
                if bin_data is not None and len(bin_data['points']) > 0:
                    points = bin_data['points']
                    x_center = x_positions[j] + offset
                    x_jitter = np.random.normal(0, bar_width/10, size=len(points))
                    x_scatter = x_center + x_jitter
                    
                    ax.scatter(x_scatter, points[fc_col], color=colors[group], edgecolor="black", linewidth=0.5,
                              alpha=0.5, s=15, zorder=20, rasterized=True)
        
        # Add lines connecting the average values for each group
        for i, group in enumerate(plot_groups):
            group_data = input_binned_df_filtered[input_binned_df_filtered['group'] == group]
            if len(group_data) > 0:
                # Create a line connecting the bar tops
                line_x = []
                line_y = []
                
                for j, bin_idx in enumerate(unique_bins):
                    bin_group_data = group_data[group_data['bin_index'] == bin_idx]
                    if len(bin_group_data) > 0:
                        # Calculate the same average used for the bars
                        if show_motifs and sequence_col in df_plot.columns:
                            bin_start = input_bins[bin_idx]
                            bin_end = input_bins[bin_idx+1]
                            
                            points_data = df_plot[(df_plot[group_col] == group) & 
                                          (df_plot[input_col] >= bin_start) & 
                                          (df_plot[input_col] < bin_end)]
                            
                            # Exclude motif points to match bar calculation
                            exclude_mask = np.zeros(len(points_data), dtype=bool)
                            if has_motif_filtered is not None:
                                exclude_mask |= has_motif_filtered.loc[points_data.index]
                            if has_custom_motif is not None:
                                exclude_mask |= has_custom_motif.loc[points_data.index]
                            
                            non_motif_points = points_data[~exclude_mask]
                            
                            if len(non_motif_points) > 0:
                                avg_fc = non_motif_points[fc_col].mean()
                            else:
                                avg_fc = 0
                        else:
                            bin_start = input_bins[bin_idx]
                            bin_end = input_bins[bin_idx+1]
                            points_data = df_plot[
                                (df_plot[group_col] == group) &
                                (df_plot[input_col] >= bin_start) &
                                (df_plot[input_col] < bin_end)]
                            if len(points_data) > 0:
                                if transform_data in ("sqrt", "log", "log_eps"):
                                # df_plot[fc_col] is already transformed
                                    avg_fc = points_data[fc_col].mean()
                                else:
                                    avg_fc = bin_group_data['avg_fc'].values[0]
                            else:
                                avg_fc = np.nan  # no bar for this bin

                            avg_fc_values.append(avg_fc)
                            bin_data_points.append({
                                'bin_index': bin_idx,
                                'x_pos': bin_idx,
                                'points': points_data
                                })
                        
                        # Position to match the bar center
                        offset = (i - len(plot_groups)/2) * bar_width
                        line_x.append(x_positions[j] + offset)
                        line_y.append(avg_fc)
                
                # Plot the connecting line
                if len(line_x) > 1:  # Only plot if we have multiple points
                    ax.plot(line_x, line_y, '-o', 
                            color=colors[group], linewidth=2.5, markersize=6, zorder=40,
                            path_effects=[path_effects.withStroke(linewidth=3, foreground='black')])
        
        # Only add motif columns if show_motifs is True
        if show_motifs:
            # Add primary motif peptides as an additional column
            if sequence_col in df_plot.columns and has_motif_filtered is not None and any(groups_to_plot):
                # Use the first group in groups_to_plot for motif plotting
                if motif_group is None:
                    motif_group = groups_to_plot[0] if groups_to_plot else 1
                if motif_group in df_plot[group_col].values:
                    print(f"\nProcessing {motif} for binned bar chart from group {motif_group}")
                    
                    motif_avg_values = []
                    motif_counts = []
                    motif_points_by_bin = []
                    
                    for bin_idx in unique_bins:
                        bin_start = input_bins[bin_idx]
                        bin_end = input_bins[bin_idx+1]
                        
                        motif_mask = (df_plot[group_col] == motif_group) & \
                                   (df_plot[input_col] >= bin_start) & \
                                   (df_plot[input_col] < bin_end) & \
                                   has_motif_filtered
                        
                        if has_custom_motif is not None:
                            motif_mask &= ~has_custom_motif
                        
                        motif_points = df_plot[motif_mask]
                        
                        if len(motif_points) > 0:
                            motif_avg = motif_points[fc_col].mean()
                            motif_avg_values.append(motif_avg)
                            motif_counts.append(len(motif_points))
                            motif_points_by_bin.append({
                                'bin_index': bin_idx,
                                'points': motif_points
                            })
                        else:
                            motif_avg_values.append(0)
                            motif_counts.append(0)
                            motif_points_by_bin.append(None)
                    
                    # Calculate the offset for the motif column
                    motif_offset = bar_width
                    
                    # Create bars for motif
                    if transform_data == "log_eps":
                        forced_ymin = -4
                        motif_bottom = forced_ymin
                        motif_heights = [v - forced_ymin for v in motif_avg_values]
                        motif_bars = ax.bar(
                            x_positions + motif_offset,
                            motif_heights,
                            width=bar_width,
                            bottom=motif_bottom,
                            #facecolor='none',
                            linewidth=0.5, 
                            edgecolor='black',
                            color='gold'
                            )
                    else:
                        motif_bars = ax.bar(
                            x_positions + motif_offset,
                            motif_avg_values,
                            width=bar_width,
                            alpha=0.8,
                            color='magenta',
                            edgecolor='black'
                            )
                    
                    # Overlay individual motif points and add connecting line
                    total_motif_points_plotted = 0
                    for j, bin_data in enumerate(motif_points_by_bin):
                        if bin_data is not None and len(bin_data['points']) > 0:
                            points = bin_data['points']
                            total_motif_points_plotted += len(points)
                            
                            x_center = x_positions[j] + motif_offset
                            x_jitter = np.random.normal(0, bar_width/10, size=len(points))
                            x_scatter = x_center + x_jitter
                            
                            ax.scatter(x_scatter, points[fc_col], 
                                      color='gold', edgecolor='black', alpha=1.0, s=30, zorder=1000, rasterized=True)
                  
                    print(f"Total {motif} points plotted in bar chart: {total_motif_points_plotted}")
                    
                    # Add connecting line
                    if any(count > 0 for count in motif_counts):
                        line_x = []
                        line_y = []
                        for j, (count, avg) in enumerate(zip(motif_counts, motif_avg_values)):
                            if count > 0:
                                line_x.append(x_positions[j] + motif_offset)
                                line_y.append(avg)
                        
                        ax.plot(line_x, line_y, '-o', 
                                color='gold', linewidth=2.5, markersize=10, zorder=30,
                                path_effects=[path_effects.withStroke(linewidth=3, foreground='black')])
        
        # Configure the plot
        ax.set_xticks(x_positions)
        ax.set_xticklabels(input_bin_labels, rotation=45, ha='right')
        ax.set_xlabel('Input Reads Bin')
        ax.set_ylabel(f'{fc_col}{y_label_suffix}')
        
        # Configure y-axis scale
        if y_scale == "log":
            if zero_count > 0:
                print("Using symlog scale to properly display zero values")
                
                linthresh = 0.01
                ax.set_yscale('symlog', linthresh=linthresh, linscale=0.2)
                
                # Use adjusted_y_max if calculated, otherwise use provided or default values
                if adjusted_y_max is not None:
                    plot_y_min = y_min if y_min is not None else fc_min
                    ax.set_ylim(plot_y_min, adjusted_y_max)
                elif y_min is not None and max_fc is not None:
                    ax.set_ylim(0, max_fc)
                else:
                    ax.set_ylim(0, fc_max)
                
                def y_fmt(x, pos):
                    if abs(x) < 0.001:
                        return '0'
                    elif x < 0.01:
                        return f'{x:.2g}'
                    elif x < 1:
                        return f'{x:.2f}'
                    else:
                        return f'{int(x)}' if abs(x - round(x)) < 1e-10 else f'{x:.1f}'
                
                ax.yaxis.set_major_formatter(ticker.FuncFormatter(y_fmt))
            else:
                ax.set_yscale('log')
                
                # Use adjusted_y_max if calculated, otherwise use provided or default values
                if adjusted_y_max is not None:
                    plot_y_min = y_min if y_min is not None else fc_min
                    ax.set_ylim(plot_y_min, adjusted_y_max)
                elif y_min is not None and max_fc is not None:
                    ax.set_ylim(y_min, max_fc)
                else:
                    ratio = fc_max / fc_min
                    if np.log10(ratio) % 2 != 0:
                        log_range = max(np.log10(fc_max), -np.log10(fc_min))
                        fc_min = 10**(-log_range)
                        fc_max = 10**log_range
                    ax.set_ylim(fc_min, fc_max)
        
        elif y_scale == "sqrt":
            print("Using square root scale for y-axis")
            
            # Handle negative values and zeros for sqrt scale
            if zero_count > 0 or (df_plot[fc_col] < 0).any():
                print("Warning: Square root scale with zero or negative values - using modified scale")
            
            # Set square root scale
            ax.set_yscale('function', functions=(lambda x: np.sqrt(np.maximum(x, 0.001)), 
                                               lambda x: x**2))
            
            # Use adjusted_y_max if calculated, otherwise use provided or default values
            if adjusted_y_max is not None:
                ax.set_ylim(max(0.001, 0), adjusted_y_max)
            elif y_min is not None and max_fc is not None:
                ax.set_ylim(max(0.001, y_min), max_fc)
            else:
                y_min_sqrt = max(0.001, df_plot[fc_col].min() * 0.9)
                ax.set_ylim(y_min_sqrt, fc_max)
            
            # Custom formatter for sqrt scale
            def sqrt_fmt(x, pos):
                if x < 0.001:
                    return '0'
                elif x < 0.01:
                    return f'{x:.3f}'
                elif x < 1:
                    return f'{x:.2f}'
                elif x < 10:
                    return f'{x:.1f}'
                else:
                    return f'{int(x)}'
            
            ax.yaxis.set_major_formatter(ticker.FuncFormatter(sqrt_fmt))
        
        else:  # linear scale
            if transform_data == "log_eps":
                # Force axis to start at -4
                forced_ymin = -4
                y_high = df_plot[fc_col].max()
                padding = 0.1 * (y_high - forced_ymin)
                ax.set_ylim(forced_ymin, y_high + padding)
            elif adjusted_y_max is not None:
                ax.set_ylim(0, adjusted_y_max)
            elif y_min is not None and max_fc is not None:
                ax.set_ylim(0, max_fc)
            else:
                if max_fc:
                    ax.set_ylim(0, max_fc)
                else:
                    ax.set_ylim(0, fc_max)
        
            
        ax.set_title(f'{fc_col} Values vs Input Reads (Binned) - Groups: {groups_to_plot}')
        ax.grid(True, alpha=1, which='both')
        ax.legend()
    
    plt.tight_layout()
    
    if output:
        plt.savefig(output)
        print(f"Saved binned bar chart to {output}")
    
    return fig, ax

def plot_logeps_fc_histograms_for_bin_by_groups(
    df,
    input_bins,
    bin_index,
    *,
    groups_to_plot,
    has_motif=None,
    motif_group=None,
    input_col="input_average",
    fc_col="FC",
    group_col="group",
    eps=1e-3,
    hist_bins=40,
    output=None
):
    """
    Vertically stacked log_eps FC histograms with KDE slope.
    One panel per group + optional motif panel.

    Parameters
    ----------
    groups_to_plot : list[int]
        Group numbers to plot (in this order)
    motif_group : int or None
        Group number from which motif peptides are taken
    """

    # -------------------------
    # 0. Defaults
    # -------------------------
    if motif_group is None:
        motif_group = groups_to_plot[0]

    # -------------------------
    # 1. Extract bin
    # -------------------------
    bin_start = input_bins[bin_index]
    bin_end = input_bins[bin_index + 1]

    bin_df = df[
        (df[input_col] >= bin_start) &
        (df[input_col] < bin_end)
    ].copy()

    if bin_df.empty:
        raise ValueError("No data in selected bin.")

    # -------------------------
    # 2. log_eps transform (same as bar chart)
    # -------------------------
    fc_vals = bin_df[fc_col].astype(float).values
    fc_vals = np.where(fc_vals < 0, 0, fc_vals)
    bin_df["log_eps_fc"] = np.log10(fc_vals + eps)

    # -------------------------
    # 3. Build datasets dynamically
    # -------------------------
    datasets = []

    for g in groups_to_plot:
        # If no motif information, just plot the group normally
        if has_motif is None or motif_group is None or g != motif_group:
            vals = bin_df.loc[
                bin_df[group_col] == g, "log_eps_fc"
            ]
            label = f"Group {g}"
        else:
            # --- EDITED ELSE STATEMENT ---
            # Now plots ALL data for this group (Total), instead of subtracting the motif
            vals = bin_df.loc[
                (bin_df[group_col] == g), 
                "log_eps_fc"
            ]
            label = f"Group {g} (Total)" 
            # -----------------------------

        if len(vals) > 0:
            datasets.append((label, vals))

    # Motif panel
    if has_motif is not None and motif_group is not None:
        motif_vals = bin_df[
            (bin_df[group_col] == motif_group) &
            (has_motif.loc[bin_df.index])
        ]["log_eps_fc"]

        if len(motif_vals) > 0:
            datasets.append((f"Motif (Group {motif_group})", motif_vals))

        if not datasets:
            raise ValueError("No valid datasets to plot.")

    # -------------------------
    # 4. Shared x bins
    # -------------------------
    all_vals = pd.concat([vals for _, vals in datasets])
    x_bins = np.linspace(all_vals.min(), all_vals.max(), hist_bins)

    # -------------------------
    # 5. Plot
    # -------------------------
    fig, axes = plt.subplots(
        nrows=len(datasets),
        ncols=1,
        figsize=(7, 2.4 * len(datasets)),
        sharex=True
    )

    if len(datasets) == 1:
        axes = [axes]

    palette = sns.color_palette("husl", len(datasets))

    x_cut = None
    for i, (ax, (label, values), color) in enumerate(zip(axes, datasets, palette)):
        values = values.dropna()

        ax.hist(
            values,
            bins=x_bins,
            density=True,
            alpha=0.6,
            color=color,
            edgecolor="black"
        )

        # KDE
        if len(values) > 5:
            sns.kdeplot(values, ax=ax, color="black", linewidth=2)

        # ---- define cutoff based on TOP plot only ----
        if i == 0:
            x_cut = np.nanquantile(values, 0.99)  # log-space cutoff

        # ---- draw same cutoff on ALL panels ----
        if x_cut is not None:
            ax.axvline(x_cut, color="black", linestyle="--", linewidth=2)

            left_count = int((values <= x_cut).sum())
            right_count = int((values > x_cut).sum())

            ax.text(
                x_cut,
                ax.get_ylim()[1] * 0.95,
                f"Left={left_count}\nRight={right_count}",
                ha="right",
                va="top",
                fontsize=9
            )

        ax.set_ylabel("Density")
        ax.set_title(f"{label} (n = {len(values)})")
        ax.grid(True, alpha=0.3)


    # --- These must be OUTSIDE the loop ---
    axes[-1].set_xlabel(r"$\log_{10}(\mathrm{FC} + \varepsilon)$")

    fig.suptitle(
        f"log_eps FC distributions\n"
        f"Input bin {bin_index + 1}: {bin_start:.0f}–{bin_end:.0f}",
        y=1.02
    )


    plt.tight_layout()

    if output:
        plt.savefig(output, dpi=300)
        print(f"Saved figure to {output}")

    return fig, axes


def quick_plot(file_path, motif=None, groups_to_plot=None, show_motifs=True, output_prefix="plot", 
               y_scale="log", transform_data="none", min_bar_height_ratio=0.33, baseline_group=None):
    """
    Quick function to generate binned bar plot with minimal setup.
    
    Parameters:
    -----------
    file_path : str
        Path to the CSV file containing peptide data
    motif : str, optional
        Motif sequence to highlight (e.g., "DMT")
    groups_to_plot : list, optional
        List of group numbers to include in the plot
    show_motifs : bool, default=True
        Whether to highlight sequences containing the motif
    output_prefix : str, default="plot"
        Prefix for output filename
    y_scale : str, default="log"
        Y-axis scale: "log", "linear", or "sqrt"
    transform_data : str, default="none"
        Data transformation: "none", "sqrt", "log"
    min_bar_height_ratio : float, default=0.33
        Position where baseline group mean should appear (0.33 = 1/3 of plot height).
        The y-axis will be automatically scaled so the mean of baseline group bars appears at this position.
        Set to 0 or None to disable automatic y-axis scaling.
    baseline_group : int, optional
        Which group number to use as baseline. If None, uses the lowest numbered group.
    
    Returns:
    --------
    matplotlib.figure.Figure
        The created bar chart figure
    """
    # Load data
    df = load_data(file_path)
    if df is None:
        return None
    
    # Check available groups
    check_available_groups(df)
    
    # Create motif mask if motif is specified
    has_motif = None
    if motif and show_motifs:
        has_motif = identify_motif_sequences(df, motif)
    
    # Process bins
    baseline_group_id = None
    if 'group' in df.columns:
        if baseline_group is not None:
            baseline_group_id = baseline_group
            print(f"Using user-specified baseline group {baseline_group_id} for tail merge")
        else:
            baseline_group_id = df['group'].min()
            print(f"Auto-detected baseline group {baseline_group_id} for tail merge")
    
    # Compute max input for baseline as auto threshold
    tail_merge_threshold = None
    if baseline_group_id is not None and 'input_average' in df.columns:
        baseline_max_input = df.loc[df['group'] == baseline_group_id, 'input_average'].max()
        tail_merge_threshold = baseline_max_input
        print(f"Tail merge threshold set to baseline max input: {baseline_max_input:.3f}")
    
    # Process bins with optional tail merge
    input_bins, input_bin_labels, input_binned_df = process_data_bins(
        df,
        tail_merge_threshold=tail_merge_threshold
    )
    
    # Create bar chart only
    fig_bar = plot_binned_bar_chart(
        df,
        input_binned_df,
        input_bins,
        input_bin_labels,
        has_motif=has_motif,
        output=f"{output_prefix}_binned.png",
        groups_to_plot=groups_to_plot,
        show_motifs=show_motifs,
        motif=motif,
        y_scale=y_scale,
        transform_data=transform_data,
        min_bar_height_ratio=min_bar_height_ratio,
        baseline_group=baseline_group
    )
    
    print(f"Bar chart saved as: {output_prefix}_binned.png")
    return fig_bar
```
