## Supplementary material for "Universal Baseline for *in vitro* Selection of Genetically Encoded Libraries": Dataset: plotting (GL 20260204).ipynb

```
# %% [markdown]
# Quick Start: DEL Bar Chart Plotting

⚡ **Fast setup guide** for creating bar charts from your DEL data.

Just update the file path and run all cells!

# %% [markdown]
## Step 1: Setup

# %% [code]
# Import libraries
%load_ext autoreload
%autoreload 2
from bin_bar_chart import quick_plot, plot_binned_bar_chart, load_data, process_data_bins, identify_motif_sequences, plot_logeps_fc_histograms_for_bin_by_groups
import matplotlib as plt
import matplotlib as plt
plt.rcParams.update({'font.size': 7, 'font.family': 'arial', 'pdf.fonttype': 42})

# Settings for better plots
plt.rcParams['figure.dpi'] = 100
plt.rcParams['savefig.dpi'] = 300

print("✅ Ready to plot!")

# %% [markdown]
## Step 2: Set Your Parameters

📝 **Edit these values** for your data:

# %% [code]
# YOUR DATA FILE
data_file = "disulfide2_KYL_reads_cpm_FC.csv"  

# YOUR SETTINGS
motif_to_highlight = "DMT"   # <-- CHANGE THIS!
baseline_group = 2     # Which group is baseline
library_group = 1      # Which group is library

# PLOT SETTINGS
baseline_position = 0.15      # 0.33 = baseline mean at 1/3 height
y_axis_scale = "log"          # "log", "linear", or "sqrt"
colors = {
    baseline_group: '#808080',  # Gray baseline
    library_group:  '#0000FF'   # Blue library
}

df = load_data(data_file)
print(f"📊 Will plot groups: {baseline_group} (baseline) and {library_group} (library)")
print(f"🎯 Baseline mean will appear at {baseline_position*100:.0f}% plot height")

# %% [code]
has_motif = identify_motif_sequences(df, motif="DMT")
input_bins, input_bin_labels, input_binned_df = process_data_bins(df)
plot_logeps_fc_histograms_with_slope_for_bin(
    df,
    input_bins,
    has_motif,
    bin_index=2,
    output="Figure1khistogram.pdf"
)
plt.show()

# %% [markdown]
## Step 3: Quick Plot (Simplest!)

One function call creates your plot:

# %% [code]
has_motif = identify_motif_sequences(df, "DMT")
baseline_group_id = 2
baseline_inputs = df.loc[df["group"] == baseline_group_id, "input_average"]
baseline_max_input = baseline_inputs.max()
print("Baseline max input:", baseline_max_input)
input_bins, input_bin_labels, input_binned_df = process_data_bins(
    df,
    input_col="input_average",
    fc_col="FC",
    group_col="group",
    tail_merge_threshold=baseline_max_input
)
print("Final bins after possible merge:", input_bins)
print("Final labels:", input_bin_labels)
# PLOT - Only change: 0.33 → 0.15
fig, ax = plot_binned_bar_chart(
    df, input_binned_df, input_bins, input_bin_labels,
    has_motif=has_motif,
    output="Figure3e.pdf",
    groups_to_plot=[baseline_group, library_group],
    colors=colors,
    baseline_group=baseline_group,
    motif_group=1,
    show_motifs=True,
    motif="DMT",
    y_scale="linear",
    transform_data="log_eps", 
    min_bar_height_ratio=0
)

plt.show()
plt.show()


# %% [code]
# YOUR DATA FILE
data_file = "disulfide2_KYL_reads_cpm_FC.csv"  

# YOUR SETTINGS
ns3_group = 3
baseline_group = 2     # Which group is baseline
chitin_group = 4      # Which group is library

# PLOT SETTINGS
baseline_position = 0.15      # 0.33 = baseline mean at 1/3 height
y_axis_scale = "log"          # "log", "linear", or "sqrt"
colors = {
    baseline_group: '#808080',  # Gray baseline
    ns3_group:  '#0000FF',# Blue library
    chitin_group: '#008000' 
}

df = load_data(data_file)


# %% [code]
groups_to_plot = [2, 1]  # [2, 3, 4] for Fig 2 and [2, 1, 0]
input_bins, input_bin_labels, input_binned_df = process_data_bins(df)
plot_logeps_fc_histograms_for_bin_by_groups(
    df,
    input_bins,
    bin_index=2,
    groups_to_plot=groups_to_plot,
    has_motif=has_motif, # comment it out if you are plotting for figure 2. 
    motif_group=1,
    output="figure2chitinhistogram.pdf"
)
plt.show()

# %% [code]
baseline_group_id = 2
baseline_inputs = df.loc[df["group"] == baseline_group_id, "input_average"]
baseline_max_input = baseline_inputs.max()
print("Baseline max input:", baseline_max_input)
input_bins, input_bin_labels, input_binned_df = process_data_bins(
    df,
    input_col="input_average",
    fc_col="FC",
    group_col="group",
    tail_merge_threshold=baseline_max_input
)
print("Final bins after possible merge:", input_bins)
print("Final labels:", input_bin_labels)
# PLOT - Only change: 0.33 → 0.15
fig, ax = plot_binned_bar_chart(
    df, input_binned_df, input_bins, input_bin_labels,
    has_motif=has_motif,
    output="ns3mix.pdf",
    groups_to_plot=[baseline_group, ns3_group, chitin_group],
    colors=colors,
    baseline_group=baseline_group,
    show_motifs=False,
    motif="XXX",
    y_scale="linear",
    transform_data="log_eps", 
    min_bar_height_ratio=0
)

plt.show()
plt.show()


# %% [code]
```
