## Supplementary material for "Universal Baseline for *in vitro* Selection of Genetically Encoded Libraries": Dataset: cleaning.py

```
import zipfile
import pandas as pd
import argparse
from data_processing import preprocess_data, aggregate_reads, merge_data, assign_group
from io import StringIO

def unzip_and_process(zip_path, ready_for_cleaning):
    print("Function called with:", zip_path, ready_for_cleaning)
    """
    Unzips the provided zip file, extracts the first .txt file, and processes it by:
    - Removing headers
    - Deleting the first 5 columns
    - Keeping columns starting after the 'AA' column

    Parameters:
    - zip_path (str): Path to the zip file.
    - output_file (str): Path to save the processed .csv file.
    """
    # Step 1: Unzip the file and locate the first .txt file
    with zipfile.ZipFile(zip_path, 'r') as zip_ref:
        print("Zip file opened successfully.")
        zip_ref.extractall("extracted_files")
        # Get the list of extracted files
        file_list = zip_ref.namelist()
        print("Extracted files:", file_list)
        # Find the first .txt file
        txt_file = next((f for f in file_list if f.endswith('.txt')), None)
        if not txt_file:
            raise FileNotFoundError("No .txt file found in the zip archive.")
        print("Processing file:", txt_file) 
        txt_path = f"extracted_files/{txt_file}"
        
# Step 2: Remove the header and extract data lines
    print("Removing header...")
    remaining_lines = []
    with open(txt_path, 'r') as file:
        for line in file:
            if line.startswith("0"):  # Adjust marker to detect data start
                remaining_lines.append(line)
                break
        for line in file:
            remaining_lines.append(line)

    print("Remaining lines after header removal:")
    print("\n".join(remaining_lines[:10]))  # Show first 10 lines

    
    # Step 3: Convert remaining lines to a DataFrame
    cleaned_data = "\n".join(remaining_lines)
    df = pd.read_csv(StringIO(cleaned_data), sep='\\s+')  # Use space delimiter
    print("DataFrame shape after loading:", df.shape)
    print("DataFrame columns:", df.columns)

    # Validate DataFrame
    if df.empty:
        raise ValueError("The DataFrame is empty. Check the input file format.")


    # Step 4: Skip the first 5 columns
    if len(df.columns) <= 5:
        raise ValueError("Not enough columns to skip the first 5.")
    print("Skipping the first 3 columns...")
    df = df.iloc[:, 3:]
    print("Columns after skipping the first 3:", df.columns)

      # Step 5: Keep the last N columns (1, 2, or 3)
    num_columns_to_keep = min(len(df.columns), 6)  # Dynamically handle up to 3 columns
    df = df.iloc[:, -num_columns_to_keep:]
    print(f"Keeping the last {num_columns_to_keep} columns:")
    print(df.head())

    
     # Rename columns dynamically
    new_column_names = ['SDB']+['DNA']+['peptide'] + [f'reads{i+1}' for i in range(num_columns_to_keep - 3)]
    print("New column names:", new_column_names)
    if len(df.columns) != len(new_column_names):
        raise ValueError(
            f"Column length mismatch: DataFrame has {len(df.columns)} columns, "
            f"but new_column_names has {len(new_column_names)}."
        )

    df.columns = new_column_names
    print("Renamed columns:", df.columns)  
    
    # Step 6: Save to CSV
    print(f"Saving to: {ready_for_cleaning}")
    df.to_csv(ready_for_cleaning, index=False)
    print("File saved successfully.")
    
if __name__ == "__main__":
    parser = argparse.ArgumentParser(description="Unzip and process a zip file containing a .txt file.")
    parser.add_argument("--zip_path", required=True, help="Path to the input zip file.")
    parser.add_argument("--ready_for_cleaning", required=True, help="Path to save the processed CSV file.")
    args = parser.parse_args()

    # Call the function with command-line arguments
    unzip_and_process(args.zip_path, args.ready_for_cleaning)

def cleaning(input_file1, input_file2, output_file, pattern=r'^S...C.........C', library="WL0.SDB21", baseline="WL0.SDB17"):
    # Load the data
    df1 = pd.read_csv(input_file1, index_col=False)
    df2 = pd.read_csv(input_file2, index_col=False)

    # Preprocess the data
    df1 = preprocess_data(df1, pattern=pattern, library=library, baseline=baseline)
    df2 = preprocess_data(df2, pattern=pattern, library=library, baseline=baseline)

    # Aggregate reads
    df1_aggregated = aggregate_reads(df1)
    df2_aggregated = aggregate_reads(df2)

    # Merge the data
    df_merged = merge_data(df1_aggregated, df2_aggregated)

    print(f"Saving merged data to {output_file}")
    df_merged.to_csv(output_file, index=False)
    print(f"File saved successfully to {output_file}")
    

if __name__ == '__main__':
    # Set up argument parser
    parser = argparse.ArgumentParser(description='Clean and merge peptide data files.')
    parser.add_argument('--cleaned', type=str, required=True, help='Path to the first input CSV file')
    parser.add_argument('--cleaned', type=str, required=True, help='Path to the second input CSV file')
    parser.add_argument('--forDE', type=str, required=True, help='Path to the output CSV file')
    parser.add_argument("--pattern", default=r'^S...C.........C', help="Regex pattern for filtering peptides.")
    # Parse arguments
    args = parser.parse_args()
    
    # Call the cleaning function with parsed arguments
    cleaning(args.input1, args.input2, args.output, pattern=args.pattern)
```
