## Supplementary material for "Universal Baseline for *in vitro* Selection of Genetically Encoded Libraries": Dataset: data_cleaning.py

```
#!/usr/bin/env python3
"""
Complete data processing script for NGS analysis pipeline
Handles dual unzipping for input and output files
"""

import argparse
import json
import os
import pandas as pd
import subprocess
import sys
from cleaning import unzip_and_process
from cleaning import cleaning
from data_processing import process_afterDE
from DE_plot import plot_input_vs_output

def parse_arguments():
    """Parse command line arguments."""
    parser = argparse.ArgumentParser(description='Process NGS data pipeline')
    parser.add_argument('--config', type=str, default='config.json',
                        help='Path to configuration file (default: config.json)')
    parser.add_argument('--skip-to-step', type=int, default=1,
                        help='Skip to specific step (1-6)')
    parser.add_argument('--run-r', action='store_true',
                        help='Run the R script for TMM normalization')
    parser.add_argument('--input-pattern', type=str, default='input',
                        help='Pattern to identify input/control columns in CSV')
    parser.add_argument('--output-pattern', type=str, default='output',
                        help='Pattern to identify output/test columns in CSV')
    parser.add_argument('--experiment', type=str,
                        help='Experiment name (e.g., R3) to update paths in the config')
    return parser.parse_args()

def check_file_exists(filepath, message=None):
    """Check if a file exists and print an error message if not."""
    if not os.path.exists(filepath):
        if message:
            print(f"Error: {message}")
        else:
            print(f"Error: File not found at {filepath}")
        return False
    return True

def main():
    # Parse command line arguments
    args = parse_arguments()
    
    # Load configuration from file
    if not check_file_exists(args.config, "Configuration file not found"):
        return 1
        
    with open(args.config, 'r') as config_file:
        config = json.load(config_file)
    
    
    # Create output directories if they don't exist
    output_dirs = [os.path.dirname(p) for p in [
        config['input_csv'],
        config['output_csv'], 
        config['output_file'],
        config['file_path'],
        config['output_path'],
        config['plot_output']
    ] if p]
    
    for directory in output_dirs:
        if directory and not os.path.exists(directory):
            os.makedirs(directory)
            print(f"Created directory: {directory}")
    
    # Step 1: Unzip and process input file
    if args.skip_to_step <= 1:
        print(f"Step 1: Processing input zip file: {config['input_zip']}")
        if check_file_exists(config['input_zip'], "Input zip file not found"):
            unzip_and_process(config['input_zip'], config['input_csv'])
            print(f"Unzipped input file to: {config['input_csv']}")
        else:
            return 1
    
    # Step 2: Unzip and process output file
    if args.skip_to_step <= 2:
        print(f"Step 2: Processing output zip file: {config['output_zip']}")
        if check_file_exists(config['output_zip'], "Output zip file not found"):
            unzip_and_process(config['output_zip'], config['output_csv'])
            print(f"Unzipped output file to: {config['output_csv']}")
        else:
            return 1
    
    # Step 3: Clean the data
    if args.skip_to_step <= 3:
        print("Step 3: Cleaning data...")
        if (check_file_exists(config['input_csv'], "Input CSV not found") and 
            check_file_exists(config['output_csv'], "Output CSV not found")):
            cleaning(
                config['input_csv'], 
                config['output_csv'], 
                config['output_file'], 
                pattern=config['pattern'],
                library=config['library'],
                baseline=config['baseline']
            )
            print(f"Cleaning complete. Output saved to {config['output_file']}")
        else:
            return 1
    
    # Step 4: TMM normalization in R
    if args.skip_to_step <= 4:
        print("Step 4: TMM normalization with R")
        if not check_file_exists(config['output_file'], "R input file not found"):
            return 1
            
        if args.run_r and 'r_script_path' in config:
            r_script = config['r_script_path']
            if not check_file_exists(r_script, "R script not found"):
                return 1
                
            print(f"Running R script: {r_script}")
            try:
                # Execute R script with the appropriate parameters
                r_executable = "C:/Program Files/R/R-4.3.2/bin/Rscript.exe"
                cmd = [
                    r_executable, r_script,
                    '-i', config['output_file'],
                    '-o', config['file_path'],
                    '--input_pattern', args.input_pattern,
                    '--output_pattern', args.output_pattern
                ]
                print("Executing:", " ".join(cmd))
                result = subprocess.run(cmd, check=True, capture_output=True, text=True)
                print(result.stdout)
                if result.stderr:
                    print("R script warnings/errors:")
                    print(result.stderr)
                print("R script completed successfully")
            except subprocess.CalledProcessError as e:
                print("Error running R script:")
                if hasattr(e, 'stdout'):
                    print(e.stdout)
                if hasattr(e, 'stderr'):
                    print(e.stderr)
                print("\nPlease run the R script manually:")
                print(f"Rscript {r_script} -i {config['output_file']} -o {config['file_path']} --input_pattern {args.input_pattern} --output_pattern {args.output_pattern}")
                return 1
        else:
            print("R script not specified or --run-r flag not used.")
            print(f"Please process {config['output_file']} with your R TMM normalization script")
            print(f"Expected output should be saved at: {config['file_path']}")
            if args.skip_to_step == 4:  # Only pause if user specifically wants to run this step
                input("Press Enter when R processing is complete...")
    
    # Step 5: Process after differential expression
    if args.skip_to_step <= 5:
        print("Step 5: Processing after DE...")
        # Check if the R output file exists
        if not check_file_exists(config['file_path'], "R output file not found. Please complete the R TMM normalization step first."):
            return 1
        
        process_afterDE(config['file_path'], config['output_path'], config['pattern'], config['inputtiter'], config['outputtiter'])
        print(f"Post-DE processing complete. Output saved to {config['output_path']}")
    
    # Step 6: Generate plots
    if args.skip_to_step <= 6:
        print("Step 6: Generating plots...")
        if not check_file_exists(config['df_path'], "Processed data file not found for plotting"):
            return 1
            
        df = pd.read_csv(config['df_path'])
        plot_input_vs_output(
            df, 
            motif=config.get('motif', 'DMT'),
            top_n=config.get('top_n', 100), 
            min_val=config.get('min_val', 1), 
            max_val=config.get('max_val', 10**6), 
            output_file=config['plot_output']
        )
        print(f"Plot generated and saved to {config['plot_output']}")
    
    print("Processing complete!")
    return 0

if __name__ == "__main__":
    sys.exit(main())
```
