## Supplementary material for "Universal Baseline for *in vitro* Selection of Genetically Encoded Libraries": Dataset: data_processing.py

```
import pandas as pd
import numpy as np
import re
def preprocess_data(df, pattern, library, baseline):
    """
    Preprocess the input DataFrame:
    - Filter library peptides using SDB and drop the ones that don't fit the pattern 
    - Filter baseline peptides using SDB and drop the ones that look like the library
    - Remove rows containing '*' or 'blank'
    - Trim peptides to the first 15 amino acids
    """
    lib_pattern = re.compile(f'.*SDB{library}.*', re.IGNORECASE)
    base_pattern = re.compile(f'.*SDB{baseline}.*', re.IGNORECASE)
    df['is_library'] = df['SDB'].astype(str).apply(lambda x: bool(lib_pattern.match(x)))
    df['is_baseline'] = df['SDB'].astype(str).apply(lambda x: bool(base_pattern.match(x)))
    print(f"Total peptides: {len(df)}")
    print(f"Library peptides (matching SDB{library}): {df['is_library'].sum()}")
    print(f"Library peptides matching pattern: {(df['is_library'] & df['peptide'].str.contains(pattern)).sum()}")
    print(f"Baseline peptides (matching SDB{baseline}): {df['is_baseline'].sum()}")
    print(f"Baseline peptides matching pattern: {(df['is_baseline'] & df['peptide'].str.contains(pattern)).sum()}")
    non_string_peptides = df[~df['peptide'].apply(lambda x: isinstance(x, str))]
    print("Peptides that are not strings (likely NaNs or floats):")
    print(non_string_peptides['peptide'].unique())
    df = df[df['peptide'].apply(lambda x: isinstance(x, str))]
    df = df[~df['peptide'].str.contains('\\*')]
    df = df[~df['peptide'].str.contains('blank')]
    df = df[df['peptide'].str.len() >= 9]
    df['c_count'] = df['peptide'].apply(lambda x: x.count('C'))

    baseline_to_remove = (
        (df["is_baseline"]) & 
        ((df['peptide'].str.len() > 9) &  # Remove if length > 9
        (df['peptide'].str.startswith('S') & (df['c_count'] == 2)))  # Remove if starts with S AND has 2 Cs
    )
    print(f"Baseline peptides to remove: {baseline_to_remove.sum()}")
    print(f"  - With length > 10: {((df['is_baseline']) & (df['peptide'].str.len() > 9)).sum()}")
    print(f"  - Starting with S and having 2 Cs: {((df['is_baseline']) & df['peptide'].str.startswith('S') & (df['c_count'] == 2)).sum()}")
    baseline_condition = (df["is_baseline"]) & ~baseline_to_remove & ~df["peptide"].str.contains(pattern)
    library_condition = (df["is_library"]) & df["peptide"].str.contains(pattern)
    df = df[library_condition | baseline_condition]
    print(f"Library peptides that will remain: {library_condition.sum()}")
    print(f"Baseline peptides that will remain: {baseline_condition.sum()}")
    df['peptide'] = df['peptide'].str[:15]
    lib_matches = df[df["is_library"] & df["peptide"].str.contains(pattern)]
    print("Matching peptides from library:")
    print(lib_matches['peptide'].head(10))
    df = df.drop(['is_library', 'is_baseline', 'c_count'], axis=1)
    return df

def aggregate_reads(df):
    """
    Aggregate reads columns by summing them up for each unique peptide.
    Dynamically handles cases where one or more of the columns are missing.
    """
    # Define the columns you want to aggregate
    columns_to_aggregate = ['reads1', 'reads2', 'reads3']

    # Determine which columns are actually present in the DataFrame
    available_columns = [col for col in columns_to_aggregate if col in df.columns]

    # If no specified columns are present, raise an error or return the original DataFrame
    if not available_columns:
        raise ValueError("None of the specified columns ('reads1', 'reads2', 'reads3') are present in the DataFrame.")

    # Perform the aggregation only on the available columns
    aggregation_dict = {col: 'sum' for col in available_columns}
    
    # Group by 'peptide' and aggregate
    return df.groupby('peptide').agg(aggregation_dict).reset_index()

def merge_data(df1, df2):
    """
    Merge two DataFrames on the 'peptide' column and fill missing values with 0.
    """
    input_suffix = "_input"
    output_suffix = "_output"
    df1_renamed = df1.copy()
    df2_renamed = df2.copy()
    for col in df1.columns:
        if col != 'peptide':
            df1_renamed = df1_renamed.rename(columns={col: f"{col}{input_suffix}"})
    
    # Rename columns in df2 with output suffix
    for col in df2.columns:
        if col != 'peptide':
            df2_renamed = df2_renamed.rename(columns={col: f"{col}{output_suffix}"})
    df_merged = pd.merge(df1_renamed, df2_renamed, on='peptide', how='outer', suffixes=('_input', '_output'))
    df_merged.fillna(0, inplace=True)
    return df_merged

# Assign groups based on patterns
def assign_group(sequence, library):
    if pd.notnull(sequence):  # Ensure the sequence is not NaN
        if pd.Series(sequence).str.contains(library, regex=True).any():
            return 1  # Group 1: library
        else:
            return 2  # Group 2: baseline
    return None  # Handle NaN values


def process_afterDE(file_path, output_path, pattern=r'^S...C.........C', inputtiter=1e10, outputtiter=1e7):
    """
    Process the input file to calculate averages, CPM, and fold changes for input and output reads.
    
    Parameters:
    - file_path (str): Path to the input tab-separated .txt file.
    - output_path (str): Path to save the processed .csv file.
    
    The function dynamically detects input and output columns by their names.
    """
    # Read the tab-separated file
    df = pd.read_csv(file_path, sep='\t')

    # Identify input and output columns dynamically
    input_columns = [col for col in df.columns if 'input' in col]
    output_columns = [col for col in df.columns if 'output' in col]

    if not input_columns or not output_columns:
        raise ValueError("Input or output columns not found in the file.")

    # Calculate average input and output reads
    df['input_average'] = df[input_columns].mean(axis=1)
    df['output_average'] = df[output_columns].mean(axis=1)


    # Adjust CPM values for plotting
    df['inputcpmplot'] = df['input_average'] + 0.1
    df['outputcpmplot'] = df['output_average'] + 0.1

    # Calculate fold change (FC)
    df['FC'] = df['output_average'] / df['input_average']
    
    # Group the peptides for plotting
    df['group'] = df['sequence'].apply(lambda x: assign_group(x, pattern))
    
    df['inputtiter'] = (df['input_average'] / df['input_average'].sum()) * inputtiter
    df['outputtiter'] = (df['output_average'] / df['output_average'].sum()) * outputtiter

    # Compute FractionalRecovery
    df = df[df['inputtiter'] != 0]
    df = df.dropna(subset=['inputtiter'])
    df['FractionalRecovery'] = np.where(
        df['outputtiter'] == 0,
        0,
        df['outputtiter'] / df['inputtiter']
    )

   
    # Save the processed DataFrame to a CSV file
    df.to_csv(output_path, index=False)

    # Display the first few rows of the processed DataFrame
    print(df.head())
```
