## Supplementary material for "Universal Baseline for *in vitro* Selection of Genetically Encoded Libraries": Dataset: DE_plot.py

```
import pandas as pd
import matplotlib.pyplot as plt
import seaborn as sns
import numpy as np

def plot_input_vs_output(df, motif='DMT', min_val=0.1, max_val=10**6, output_file='R3plot.png'):
    """
    Plots a log scale scatter plot of Input vs Output ppm with highlighted top N peptides and motif matches.
    
    Parameters:
    - df: pandas DataFrame containing the data with columns 'inputppmplot', 'outputppmplot', and 'sequence'.
    - top_n: int, the number of top entries to highlight (default is 100).
    - motif: str, the peptide sequence motif to highlight (default is 'DMT').
    - min_val: int, the minimum value for the diagonal line (default is 1).
    - max_val: int, the maximum value for the diagonal line (default is 10**6).
    - output_file: str, the file name for saving the plot (default is 'R2plot.png').
    """
    
    # Filter rows that contain the motif
    motif_rows = df[(df['sequence'].str.contains(motif, case=False, na=False)) & (df['group'] == 1)]
    
   
    # Create diagonal line points
    line_points = np.linspace(min_val, max_val, 100)
    
    # Create the plot
    plt.figure(figsize=(8, 8))
    
    if 'group' in df.columns:  # If 'group' column exists
        # Plot Group 2 as green
        group_baseline_df = df[df['group'].isin([2])]       # Plot other groups with a default color
        data = df[df['group'] == 1]
        plt.scatter(data['inputcpmplot'], data['outputcpmplot'], color='green', linewidth=0.2, s=80)        
        plt.scatter(group_baseline_df['inputcpmplot'], group_baseline_df['outputcpmplot'], color='gray', alpha=0.25, linewidth=0.2, s=80, label='baseline')
        
    else:  # If 'group' column doesn't exist
        plt.scatter(df['inputcpmplot'], df['outputcpmplot'], color='green', linewidth=0.2, s=80, label='All Data')

    
    # Highlight the motif-containing rows in gold
    sns.scatterplot(x='inputcpmplot', y='outputcpmplot', data=motif_rows, color='magenta', edgecolor='black', s=80, linewidth=0.1, label=f'{motif} Motif')
    
    # Plot the diagonal reference line
    plt.plot(line_points, line_points, color='red', linestyle='--')
    
    # Set log scale
    plt.xscale('log')
    plt.yscale('log')
    
    # Ensure the aspect ratio is equal
    plt.gca().set_aspect('equal', adjustable='box')
    
    # Set plot labels and title
    plt.title('Input vs Output ppm (Log Scale)')
    plt.xlabel('Input (log scale)')
    plt.ylabel('Output (log scale)')
    
    # Show legend
    plt.legend()
    
    # Save the plot to a file
    plt.savefig(output_file, dpi=300, bbox_inches='tight')
    
    # Display the plot
    plt.show()
```
