## Supplementary material for "Universal Baseline for *in vitro* Selection of Genetically Encoded Libraries": Dataset: generate_config.py

```
#!/usr/bin/env python3
"""
Generate configuration files for NGS data processing pipeline
"""

import argparse
import json
import os

def parse_arguments():
    parser = argparse.ArgumentParser(description='Generate configuration for NGS data processing')
    parser.add_argument('--experiment', type=str, required=True,
                      help='Experiment name (e.g., R3)')
    parser.add_argument('--input-dir', type=str, required=True,
                      help='Directory containing input and output zip files')
    parser.add_argument('--output-dir', type=str, required=True,
                      help='Directory for final output files')
    parser.add_argument('--pattern', type=str, default='^S...C.........C',
                      help='Regex pattern for sequence filtering')
    parser.add_argument('--output-file', type=str, default='config.json',
                      help='Output config file path')
    parser.add_argument('--library', type=str, default='EJ0.SDB21',
                      help='Library identifier')
    parser.add_argument('--baseline', type=str, default='EJ0.SDB17',
                      help='Baseline identifier')
    parser.add_argument('--motif', type=str, default='DMT',
                      help='Motif for plot generation')
    parser.add_argument('--inputtiter', type=float, default=9.75e8,
                      help='Input titer value')
    parser.add_argument('--outputtiter', type=float, default=5.43e6,
                      help='Output titer value')
    return parser.parse_args()

def main():
    args = parse_arguments()
    
    # Create configuration
    config = {
        "input_zip": os.path.join(args.input_dir, "input.zip"),
        "output_zip": os.path.join(args.input_dir, "output.zip"),
        "input_csv": os.path.join(args.input_dir, f"{args.experiment}input.csv"),
        "output_csv": os.path.join(args.input_dir, f"{args.experiment}output.csv"),
        "output_file": os.path.join(args.input_dir, f"{args.experiment}forDE.csv"),
        "pattern": args.pattern,
        "library": "EJ0.SDB21",
        "baseline": "EJ0.SDB17",
        "r_script_path": "./tmm_normalization.R",
        "file_path": os.path.join(args.output_dir, f"{args.experiment}DE.txt"),
        "inputtiter": 9.75e8,
        "outputtiter": 5.43e6,
        "output_path": os.path.join(args.output_dir, "reads_cpm_FC.csv"),
        "df_path": os.path.join(args.output_dir, "reads_cpm_FC.csv"),
        "plot_output": os.path.join(args.output_dir, f"{args.experiment}plot.png"),
        "motif": "DMT",
        "top_n": 100,
        "min_val": 1,
        "max_val": 1000000
    }
    
    # Write configuration to file
    with open(args.output_file, 'w') as f:
        json.dump(config, f, indent=4)
    
    print(f"Configuration created at {args.output_file}")
    print("\nExample usage:")
    print(f"python data_processing_script.py --config {args.output_file} --run-r")

if __name__ == "__main__":
    main()
```
