## Supplementary material for "Universal Baseline for *in vitro* Selection of Genetically Encoded Libraries": Dataset: readme.md

# NGS Data Processing Package - User Guide

This guide will walk you through how to use the NGS data processing package (file downloaded from https://48hd.cloud/) for biopanning using p3 library step by step, even if you have no experience with Python.

**Important Note:** This package can be used for data processing with:
- A single library configuration, or
- Two libraries with separate SDB codes

Please download the appropriate version according to your research needs.

## Prerequisites

- Windows computer
- R installed (version 4.3.2 or higher)
- Python installed (version 3.6 or higher)

## Running the Analysis

### Step 1: Generate Configuration File

Run the following command to create a configuration file for your experiment:

```
python generate_config.py --experiment YOUR_EXPERIMENT_NAME --input-dir C:\path\to\your\zip\files --output-dir C:\path\to\desired\output\folder --pattern your_library_pattern --library SDB_for_the_library --baseline SDB_for_the_baseline
```

### Step 2: Run the Data Processing Script

```
python data_cleaning.py --config config.json --run-r --experiment YOUR_EXPERIMENT_NAME
```


## Troubleshooting

### Error: File not found

Make sure your zip files are in the correct location and named correctly (`input.zip` and `output.zip`).

### R Script Issues

If the R script fails to run:
1. Make sure R is installed correctly
2. The path to Rscript.exe might need to be updated in data_cleaning.py
3. Try running the R script manually as suggested in the error message

## Advanced Options

### Skipping Steps

If you want to run only specific steps:
```
python data_cleaning.py --config config.json --skip-to-step NUMBER
```
Where NUMBER is:
1. Process input zip (unzip the input file to txt)
2. Process output zip (unzip output file to txt)
3. Clean data (remove repetitive lines, blank and peptides that do not belong to the library, combine input and output files)
4. TMM normalization (normalize input and output)
5. Post-differential expression processing (calculate the fold change, counts per million value for plotting)
6. Generate plots 

### Changing Parameters

To modify settings like pattern matching or motifs, edit the config.json file after it's generated or update the generate_config.py script.

## Example Workflow

1. Create experiment folder: `C:\NGS_Analysis\R3_Test`
2. Copy input.zip and output.zip to that folder
3. Generate config:
   ```
   python generate_config.py --experiment R3 --input-dir C:\NGS_Analysis\R3_Test --output-dir C:\NGS_Analysis\Results\R3
   ```
4. Run the analysis:
   ```
   python data_cleaning.py --config config.json --run-r
   ```
5. View the results in the output directory
