## Supplementary material for "Universal Baseline for *in vitro* Selection of Genetically Encoded Libraries": Dataset: tmm_normalization.R

```
#!/usr/bin/env Rscript
# Automated TMM normalization script for NGS analysis
# Automatically identifies test (output) and control (input) columns

# Loading Required Libraries
suppressPackageStartupMessages({
  library(edgeR)
  library(ggplot2)
  library(knitr)
  library(optparse)
})

# Command line argument parsing
option_list = list(
  make_option(c("-i", "--input"), type="character", default=NULL, 
              help="Input CSV file path", metavar="FILE"),
  make_option(c("-o", "--output"), type="character", default=NULL, 
              help="Output file path for TMM normalized data", metavar="FILE"),
  make_option(c("--input_pattern"), type="character", default="input", 
              help="Pattern to identify control columns [default= %default]"),
  make_option(c("--output_pattern"), type="character", default="output", 
              help="Pattern to identify test columns [default= %default]")
)

opt_parser = OptionParser(option_list=option_list)
opt = parse_args(opt_parser)

# Check if required arguments are provided
if (is.null(opt$input) || is.null(opt$output)) {
  cat("Error: Input and output file paths are required\n")
  cat("Usage: Rscript tmm_normalization.R -i input_file.csv -o output_file.txt\n")
  quit(status=1)
}

# Print configuration
cat("Configuration:\n")
cat(paste("Input file:", opt$input, "\n"))
cat(paste("Output file:", opt$output, "\n"))
cat(paste("Input column pattern:", opt$input_pattern, "\n"))
cat(paste("Output column pattern:", opt$output_pattern, "\n"))

# Load and prepare the dataset
cat("Loading data...\n")
counts <- try(read.table(opt$input, header = TRUE, sep = ",", quote = "\"", check.names = FALSE))
if (class(counts) == "try-error") {
  cat("Error: Failed to read input file. Please check the file path and format.\n")
  quit(status=1)
}

# Identify control and test columns
colnames <- colnames(counts)
if (length(colnames) <= 1) {
  cat("Error: Input file must have at least one data column plus a sequence column.\n")
  quit(status=1)
}

# Find column indices (skipping the first column which should be sequence)
control_indices <- grep(tolower(opt$input_pattern), tolower(colnames[-1]), value = FALSE)
test_indices <- grep(tolower(opt$output_pattern), tolower(colnames[-1]), value = FALSE)

# Adjust indices to account for the first column being removed later
control_indices <- control_indices + 1
test_indices <- test_indices + 1

if (length(control_indices) == 0) {
  cat("Warning: No control columns found using pattern '", opt$input_pattern, "'\n")
  cat("Available columns:", paste(colnames[-1], collapse=", "), "\n")
  quit(status=1)
}

if (length(test_indices) == 0) {
  cat("Warning: No test columns found using pattern '", opt$output_pattern, "'\n")
  cat("Available columns:", paste(colnames[-1], collapse=", "), "\n")
  quit(status=1)
}

cat("Identified columns:\n")
cat("Control columns:", paste(colnames[control_indices], collapse=", "), "\n")
cat("Test columns:", paste(colnames[test_indices], collapse=", "), "\n")

# Create the group factor
all_cols <- c(control_indices, test_indices)
group <- factor(c(rep("control", length(control_indices)), 
                 rep("test", length(test_indices))))

# Make sure we only use the identified columns
counts_subset <- counts[, c(1, all_cols)]

# Assign sequence to row names
row.names(counts_subset) <- counts_subset[,1]
counts_subset <- counts_subset[,-1]

# Differential Expression Analysis using edgeR
cat("Performing TMM normalization...\n")
counts_subset[is.na(counts_subset)] <- 0
RG <- DGEList(counts = counts_subset, group = group)
RG <- calcNormFactors(RG)

tmm_data <- cpm(RG, normalized.lib.sizes = TRUE)
final_table <- data.frame(sequence = rownames(RG$counts), tmm_data)

# Write the results
cat("Writing results to", opt$output, "\n")
write.table(final_table, opt$output, quote = FALSE, sep = "\t", row.names = FALSE)
cat("TMM normalization complete!\n")
```
