## Supplementary material for "Universal Baseline for *in vitro* Selection of Genetically Encoded Libraries": Dataset: scatterplot.ipynb

```
# %% [code]
from DE_plot import plot_input_vs_output
import pandas as pd
import matplotlib.pyplot as plt
df = pd.read_csv('./chitin/S2C8C2/R1R2R3unmodified_20241219/reads_cpm_FC.csv')
plot_input_vs_output(df, motif="HPV", output_file="./chitin/S2C8C2/R1R2R3unmodified_20241219/R3plot.png")
plot_input_vs_output(df, motif="XXX", output_file="./chitin/S2C8C2/R1R2R3unmodified_20241219/R3plotnomotif.png")

# %% [code]
```
