## Supplementary figures and images for "Universal Baseline for *in vitro* Selection of Genetically Encoded Libraries"

### ns3control.png

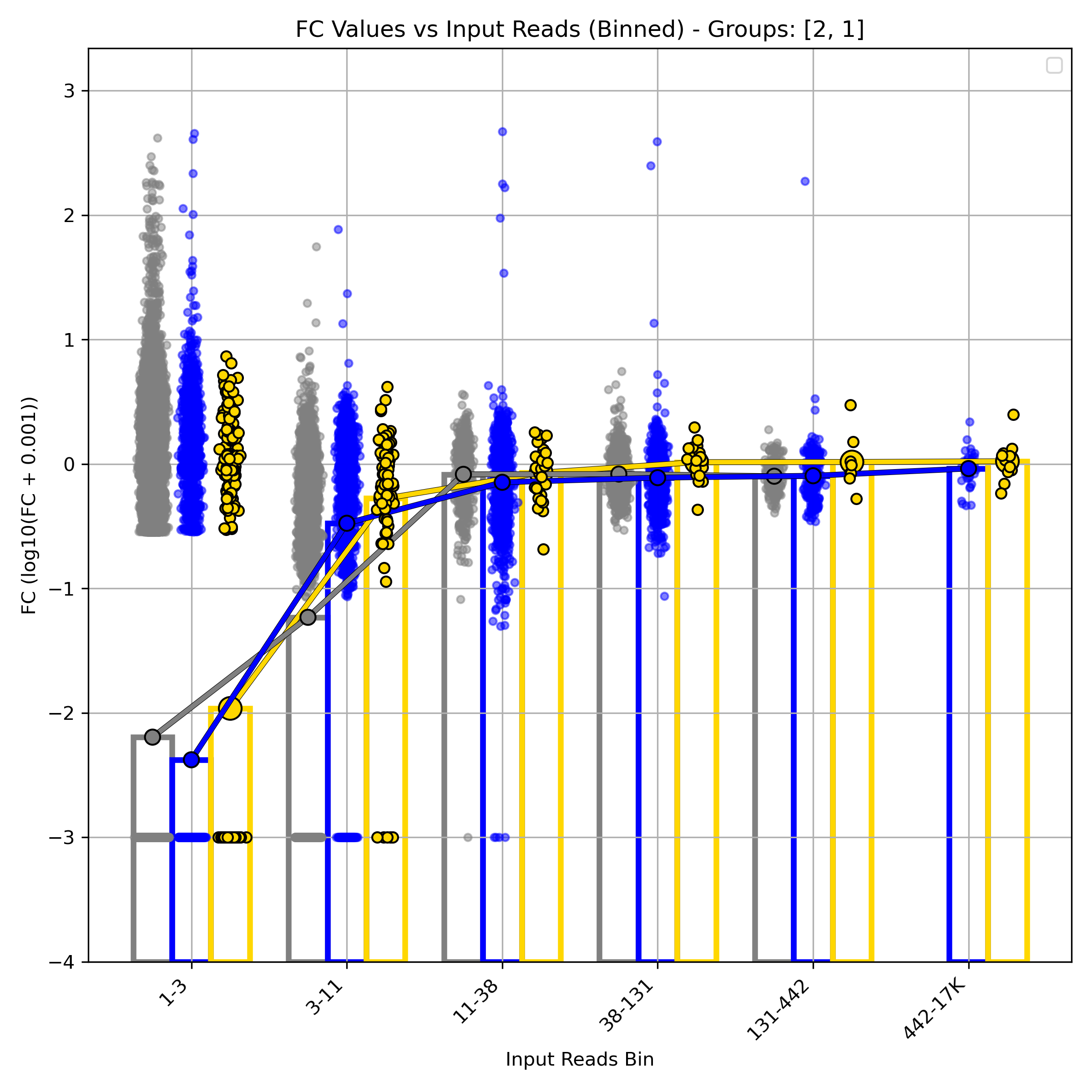

### ns3histograms.png

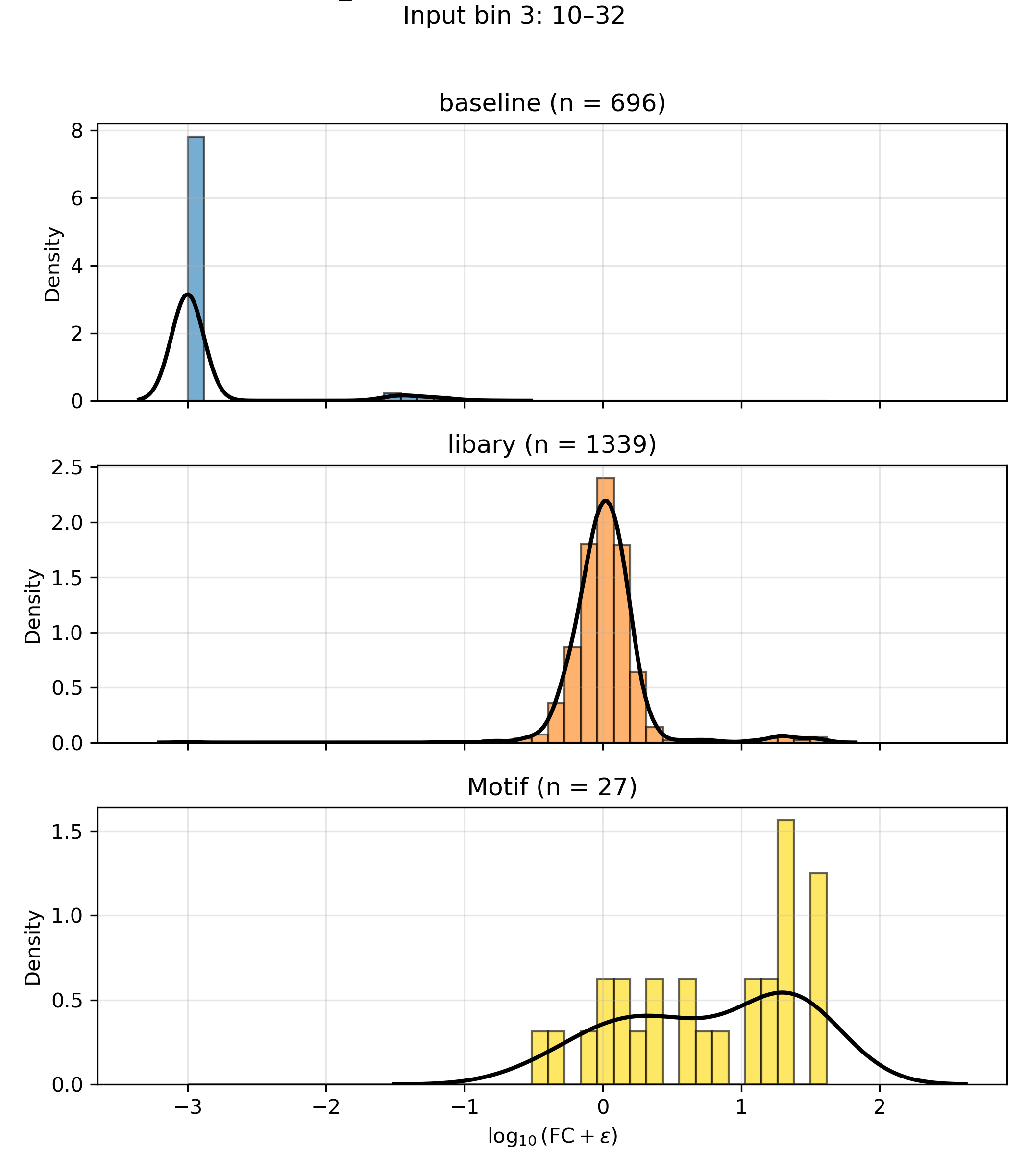
